# Cell type-specific and disease-associated eQTL in the human lung

**DOI:** 10.1101/2023.03.17.533161

**Authors:** Heini M Natri, Christina B Del Azodi, Lance Peter, Chase J Taylor, Sagrika Chugh, Robert Kendle, Mei-i Chung, David K Flaherty, Brittany K Matlock, Carla L Calvi, Timothy S Blackwell, Lorraine B Ware, Matthew Bacchetta, Rajat Walia, Ciara M Shaver, Jonathan A Kropski, Davis J McCarthy, Nicholas E Banovich

## Abstract

Common genetic variants confer substantial risk for chronic lung diseases, including pulmonary fibrosis (PF). Defining the genetic control of gene expression in a cell-type-specific and context-dependent manner is critical for understanding the mechanisms through which genetic variation influences complex traits and disease pathobiology. To this end, we performed single-cell RNA-sequencing of lung tissue from 67 PF and 49 unaffected donors. Employing a pseudo-bulk approach, we mapped expression quantitative trait loci (eQTL) across 38 cell types, observing both shared and cell type-specific regulatory effects. Further, we identified disease-interaction eQTL and demonstrated that this class of associations is more likely to be cell-type specific and linked to cellular dysregulation in PF. Finally, we connected PF risk variants to their regulatory targets in disease-relevant cell types. These results indicate that cellular context determines the impact of genetic variation on gene expression, and implicates context-specific eQTL as key regulators of lung homeostasis and disease.

## Introduction

Genomic and functional studies have the potential to reveal the genetic, molecular, and cellular drivers of clinical phenotypes, laying the groundwork for the development of targeted interventions. Many disease-associated variants identified in genome-wide association studies (GWAS) are located in the regulatory regions of the genome and contribute to disease risk and progression by effecting changes in gene expression.^1^ Combining genotype information with transcriptional profiles allows for the identification of genetic regulators of gene expression (expression quantitative trait loci, eQTL). This approach has been widely applied to bulk RNA sequencing of primary tissues, providing insights into the tissue-specificity of regulatory effects and contributing to our understanding of the mechanisms underlying complex traits.^2^ However, cell type and context (e.g., disease-status) specificity of trait-associated single-nucleotide polymorphisms (SNPs) poses a challenge to understanding the regulatory mechanisms that modulate disease risk and progression.

Single-cell RNA sequencing (scRNA-seq) has emerged as a powerful tool for the transcriptional profiling of individual cells and cell types, mitigating many limitations of bulk RNA-seq. Capturing scRNA-seq profiles and genome-wide genotype information for a population of individuals allows for the unbiased, cell type-specific interrogation of variant effects on gene expression. This approach can enable the discovery of eQTL that are specific to rare or disease-relevant cell types and eQTL that have opposing effects in different cell types, all of which could go undetected with bulk RNA-seq in heterogeneous tissues. These context-specific eQTL are thought to be more likely to escape the purifying selection that limits mutations impacting ubiquitous eQTL and are thus more likely to play roles in disease.^3,4^

Interstitial lung diseases (ILDs) are chronic, progressive respiratory disorders characterized by the scarring of lung tissue accompanied by epithelial remodeling, loss of functional lung alveoli, and accumulation of extracellular matrix (ECM).^5^ Pulmonary fibrosis (PF) is the end-stage clinical phenotype of ILD. PF remains incurable, and the most severe form of PF (idiopathic PF, IPF), leads to death or lung transplant within 3 to 5 years of diagnosis.^5,6^ The pathogenesis and progression of IPF involve a complex interplay of predisposing factors, cell types, and regulatory pathways.^7,8^ GWAS and meta-analyses have identified 20 IPF-associated variants, and polygenic analyses suggest that a large number of unreported variants contribute to IPF susceptibility.^9^ Some of these variants are eQTL in bulk-lung tissue; however, their cell type-specific regulatory consequences have not been explored.

To investigate the genetic control of disease-related gene expression in PF, we generated scRNA-seq data from lung tissue samples of 116 individuals (67 ILD and 49 unaffected donors). Combining these data with genome-wide genotype data, we mapped shared, lineage-specific, and cell type-specific *cis*-eQTL across 38 cell types (**Fig. 1a**). We analyzed these data in conjunction with IPF and other GWAS summary statistics to uncover the regulatory mechanisms underlying ILD risk and progression. Using interaction models, we reveal disease-specific regulatory effects that further elucidate the mechanisms underlying disease biology.

**Fig. 1:**
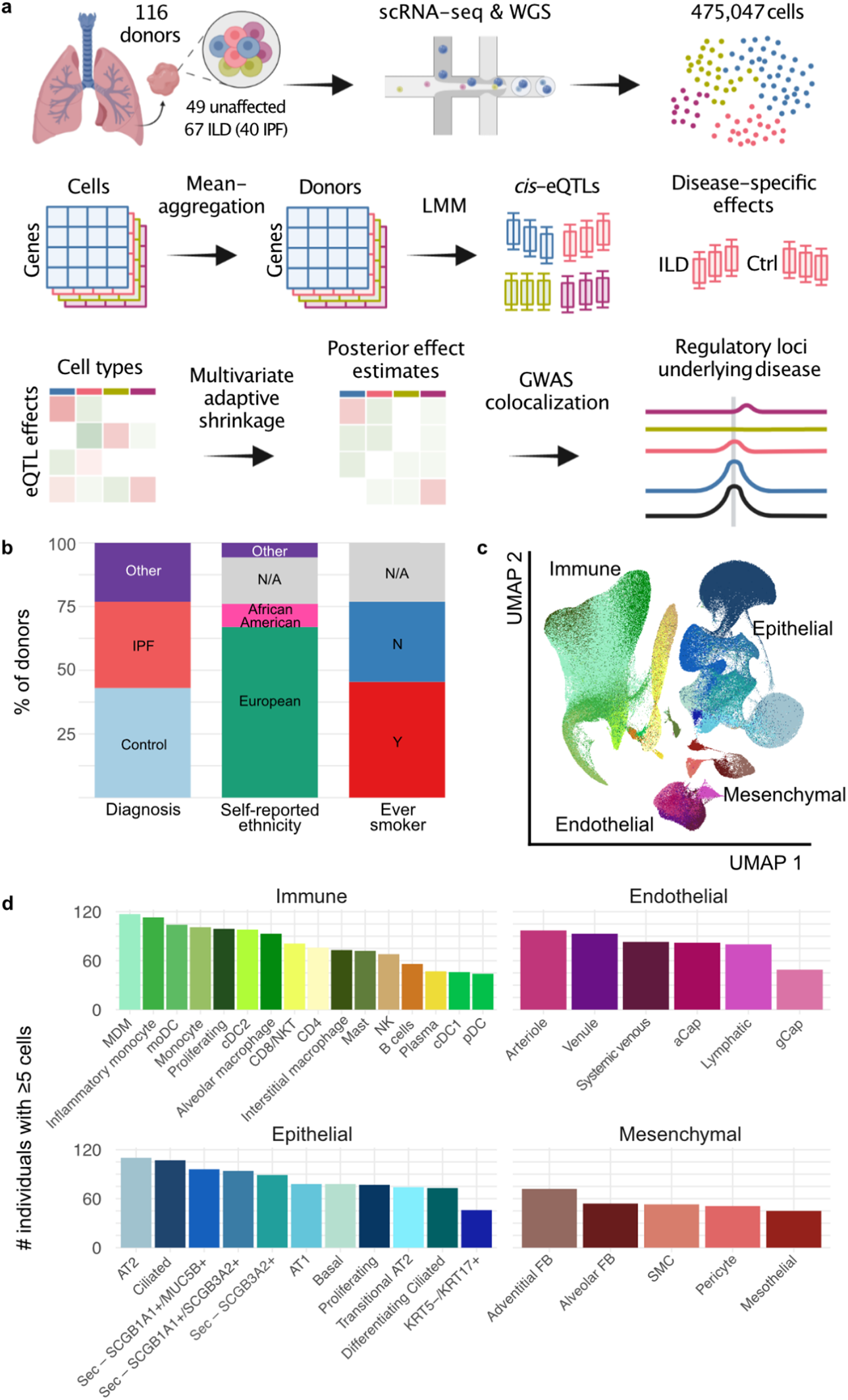
Mapping eQTL across cell types in the human lung. **a**, A schematic illustration of the present study. **b**, Percentage proportions of donors by diagnosis, self-reported ethnicity, and smoking history. **c**, a UMAP dimensionality reduction of 437,618 cells across the 38 cell types included in the eQTL analysis. Pseudocoloring indicates cell type, primary cell lineages are labeled. **d**, Numbers of donors with ≥5 cells for each cell type included in the analysis.

## Results

### Single-cell RNA-sequencing of 116 lung tissue samples

To enable cell type level eQTL mapping, we generated scRNA-seq and genome-wide genotype profiles for 116 individuals, including 67 (58%) with ILD and 49 (42%) unaffected donors (**Fig. 1a, Supplementary Table 1**). The ILD lungs included samples from 40 individuals with IPF and 27 with other forms of PF, including sarcoidosis (n=4), connective tissue disease-associated interstitial lung disease (CTD-ILD, n=3), idiopathic nonspecific interstitial pneumonia (NSIP, n=3), coal worker’s pneumoconiosis (CWP, n=3), chronic hypersensitivity pneumonitis (cHP, n=2), interstitial pneumonia with autoimmune features (IPAF, n=2), and unclassifiable ILD (n=10). The majority (68%) of the lung samples were from individuals with self-reported ethnicity information of European ancestry, and 55 (47%) reported past or present tobacco use (**Fig. 1b**).

Single-cell suspensions were generated from fresh peripheral lung tissue samples and sequenced using the 10x Genomics Chromium platform. For 55 ILD lungs, two libraries were prepared from differentially involved areas of one lung to account for regional heterogeneity. Genotype data was obtained through low pass whole genome sequencing followed by imputation (**Methods**). We performed data integration, dimensionality reduction and unsupervised clustering of the 475,047 cells passing QC using the Seurat package^10^ (**Methods**). Based on marker gene expression (**Supplementary Table 2**), we identified 43 cell types with a median of 5,811 cells (min=253, max=94,413, mean=11,048) cells (**Fig. 1c**).

### Most eQTL are shared between cell types

Out of 43 annotated cell types, we selected 38 that had ≥40 donors with ≥5 cells for that cell type to use for eQTL discovery (**Fig. 1d**). Pseudo-bulk eQTL mapping was performed on each cell type using LIMIX following the optimized approach described in Cuomo, Alvari, Azodi et al^11^. To maximize precision and overcome varying statistical power across cell types, we used multivariate adaptive shrinkage, a statistical method for analyzing measures of effect sizes across many conditions to identify patterns of sharing and specificity^12^. After applying multivariate adaptive shrinkage with mashr (**Methods**), eQTL were considered significant if they had a local false sign rate (lfsr) ≤0.05 in at least one cell type and ≤0.1 in any additional cell types. A gene was considered an eGene for a cell type if any eQTL for that gene was significant. Of the 6,995 genes tested for eQTL (**Methods**), 6,637 (94%) were eGenes in at least one cell type. The number of eGenes found per cell type was greater for more abundant cell types (**Fig. 2a**), with a positive correlation (R=0.66, *p*=6.6×10^−6^) between the number of eGenes and the number of individuals used for mapping (**Fig. 2b**).

**Fig. 2:**
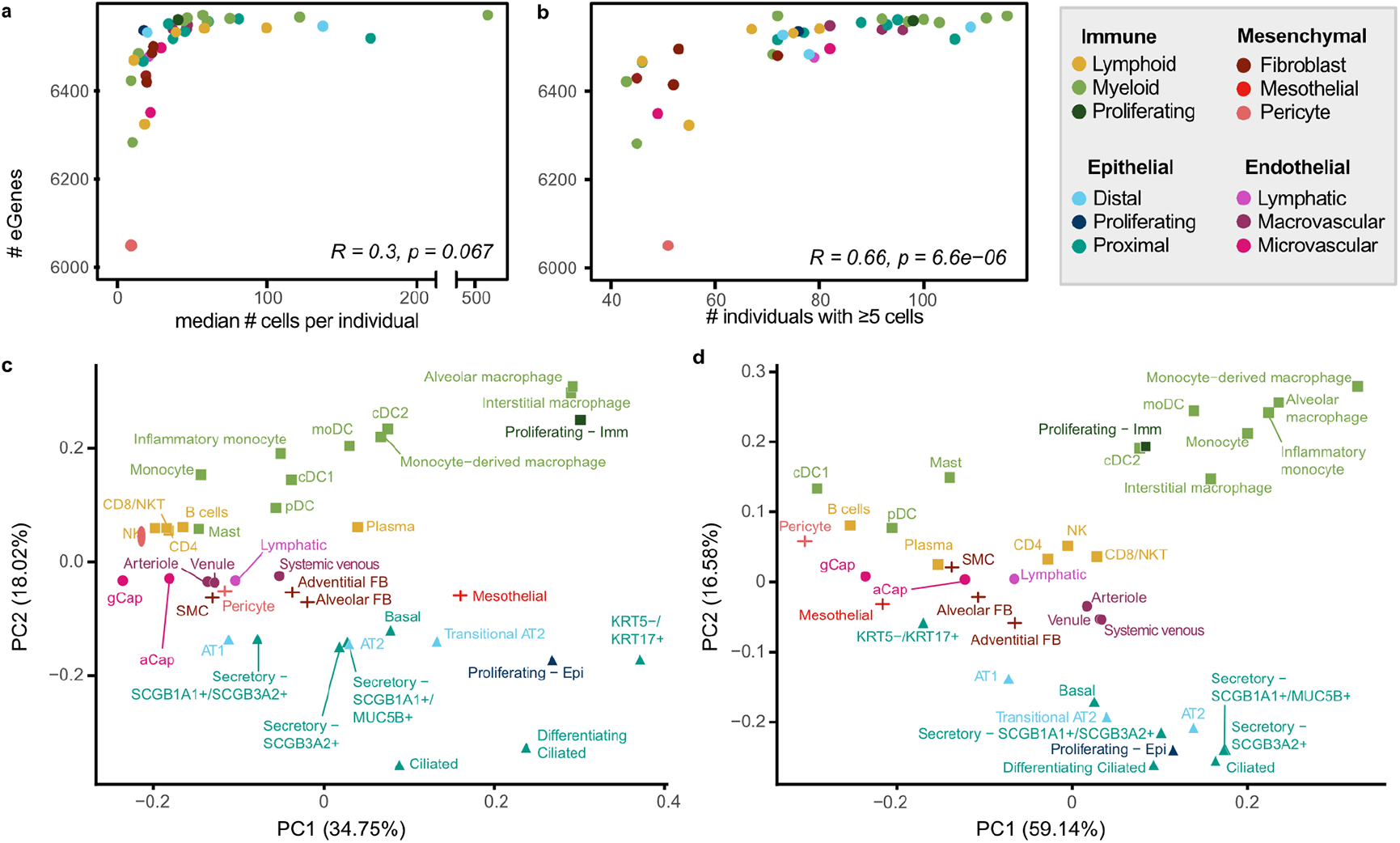
sc-eQTL reflect lineage and cell type relationships of expression measurements. **a**, Comparison of the number of eGenes per cell type and the median number of cells per individual of that cell type (Pearson’s correlation). Cell types are colored by sublineage. **b**, Comparison of the number of eGenes per cell type and the number of individuals with at least 5 cells of that cell type (Pearson’s correlation). **c**, PCA plot of pseudo-bulk expression across the 6,995 genes included in the eQTL mapping analysis. **d**, PCA plot of mashr estimated effect sizes for top eQTL (n=50,389).

To summarize the overall pattern of eQTL sharing between cell types and compare this pattern with the transcriptional similarity, we visualized the top two principal components of the median pseudo-bulked gene expression levels across all 38 cell types for the 6,995 genes included in eQTL mapping (**Fig. 2c**) and of the mashr estimated effect sizes of top eQTL across all 38 cell types (**Fig. 2d**). This demonstrated that the relationships between the regulatory mechanisms across lung cell types largely reflected the differences in expression patterns across cell types. We identified a set of top-eQTL by selecting the eQTL with the lowest, significant lfsr for each gene in each cell type. Using this criteria, there were 50,389 top eQTL, with a median of 7 top eQTL per gene (min=1, max=33). Top eQTL are considered shared between two cell types if they are significant in both cell types and their mashr estimated effect size is within a factor of 0.5. Across all cell types, the median pairwise sharing of top eQTL was 93.5% (min=55%, max=99.3%; **Fig. 3**). The epithelial and endothelial lineages had the highest levels of inter-lineage sharing (median=97.9%), while sharing between cell types within the mesenchymal lineage (median=96.9%) and the immune lineages (median=95.4%) was slightly lower.

**Fig. 3:**
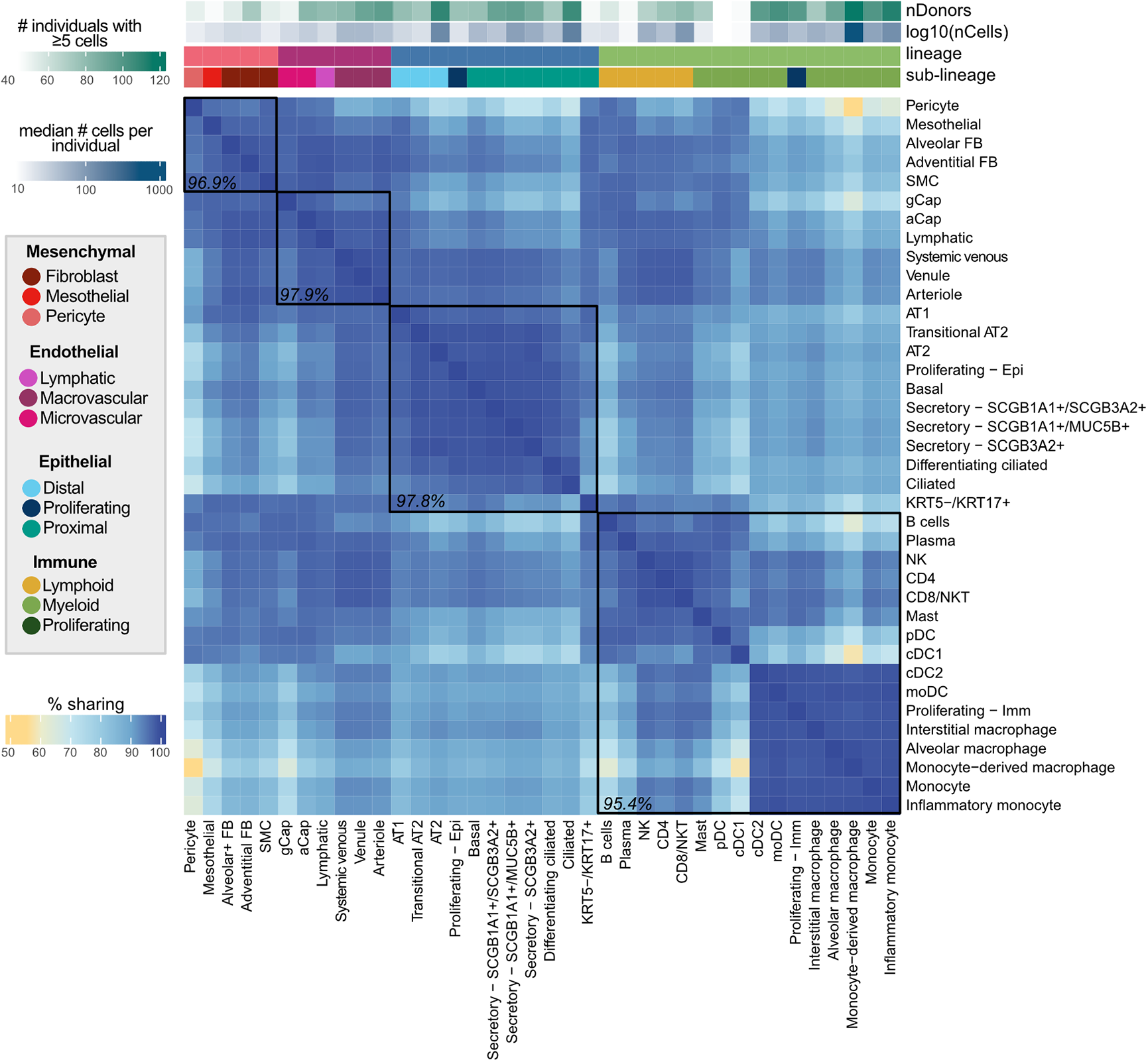
eQTL are largely shared between lung cell types. Percent of top eQTL (n=50,389) that are shared between two cell types. Top eQTL are considered shared if they are significant in both cell types (local false sign rate ≤0.1) and the mashr estimated effect size is within a factor of 0.5. Cell types are annotated above by lineage, sublineage, the number of individuals with ≥5 cells, and the median number of cells per individual for that cell type. Median pairwise percent sharing per lineage is shown in black.

We further classified top-eQTL as global (n=34,030), multi-cell-type (n=14,027), or unique to a specific cell type (n=2,332) (**Methods**). Global top-eQTL tended to be found for genes that had higher average expression and that were more widely expressed across cells (**Fig. 4a**). Top-eQTL unique to a single cell type tended to have higher absolute estimated effect sizes, likely due to limited statistical power to detect cell type-specific effects (**Fig. 4a**). Finally, these unique top-eQTL also tended to be located further from the transcription start site (TSS; **Fig. 4a**) of their target, consistent with the observation that cell type-specific eQTL typically impact enhancers, while widely shared eQTL impact promoters.^13,14^

**Fig. 4:**
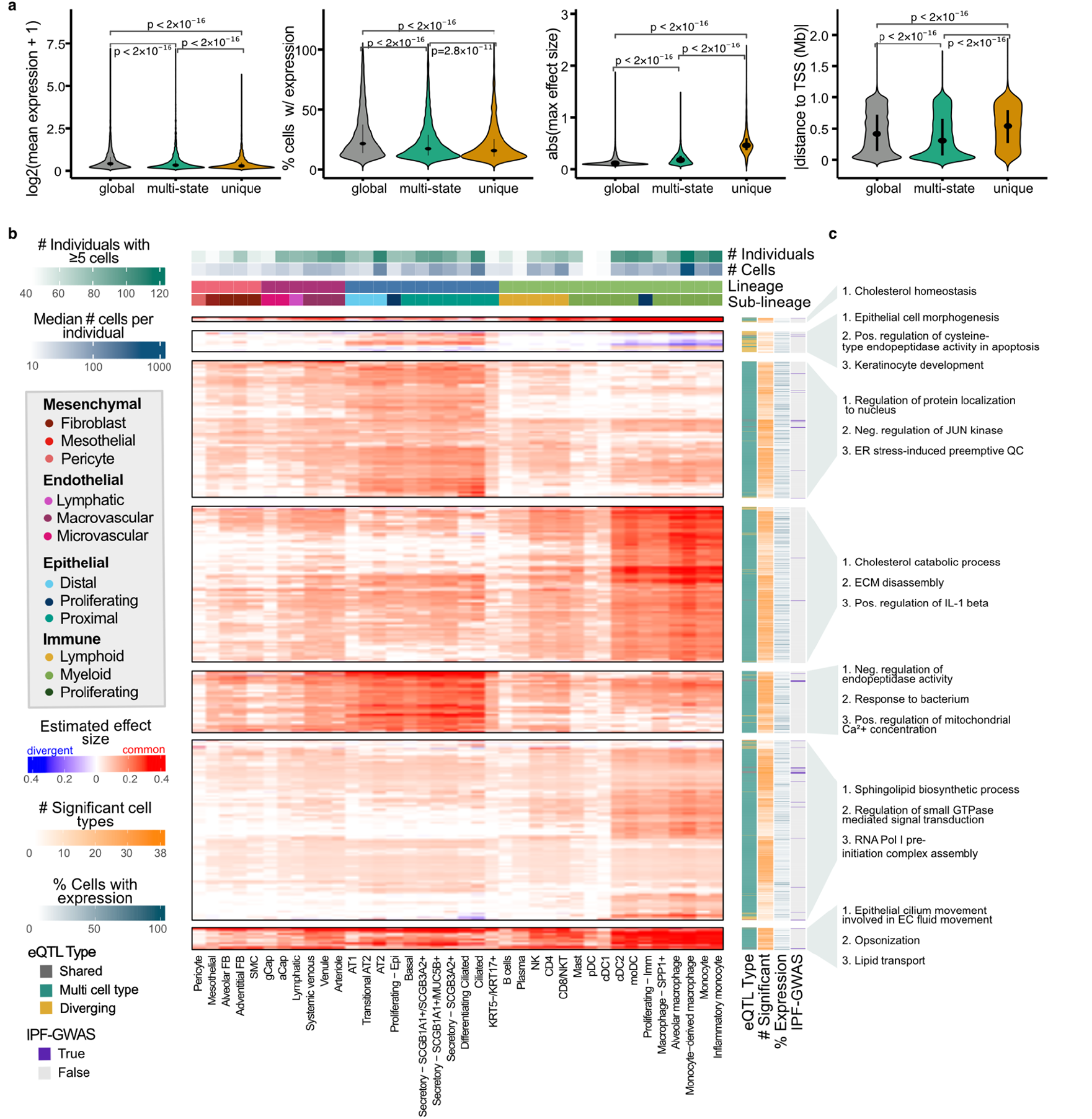
Multi-cell type eQTL act in a highly lineage-specific manner. **a**, Mean normalized expression (first panel) and percentage proportion of cells expressing (second panel) the target eGenes of eQTL unique to a single cell type, shared across multiple cell types, or globally shared across all cell types, as well as the absolute effect sizes (third panel) and absolute distances to the target eGene TSS (fourth panel). **b**, Visualization of a representative subset (**Methods**) of multi-cell-type top-eQTL and IPF-GWAS eQTL (n=3,725). eQTL are clustered by their estimated effect sizes, with non-significant associations set to zero. The most common effect direction for each eQTL is shown in red and cell types with opposite effect directions are shown in blue. **c**, Top three most significantly enriched gene ontology terms for each cluster in **b**, excluding terms with support from <2 genes.

To explore the pattern of eQTL sharing across cell types more closely we focused on two sets of top-eQTL: (1) those with known associations to IPF and (2) multi-cell-type top-eQTL. Top-eQTL were considered to be associated with IPF if the identified eGene was previously reported (*p*<1×10^−12^ in an IPF GWAS meta-analysis^9^). We further pruned these top-eQTL to get a representative sample for plotting (n=3,725) and adjusted the sign of the effect sizes to where positive indicates the common effect direction and negative indicates an opposite effect direction; **Methods**). In an unsupervised clustering of the sign-adjusted effect sizes of these pruned eQTL, we identify distinct classes of eQTL (**Fig. 4b;** k1-k7 top to bottom), including groups of eQTL primarily active in epithelial or immune cell types, or exhibiting opposing effects between lineages. To connect these eQTL to biological processes, we tested for the enrichment of their target eGenes among Gene Ontology terms against a set of 6,995 background genes (**Fig. 4c, Methods**). The eQTL in cluster 3 were primarily active in the epithelial cell types and were enriched for genes involved in the regulation of JUN kinase, which has been implicated in lung fibrosis and is a potential target for interventions for ILD.^15^ Epithelial eQTL in cluster 5 were enriched for genes associated with metabolism and response to bacteria. The eQTL in cluster 4 were primarily significant in the myeloid innate immune cell types, and showed enrichment for genes involved in, e.g., cholesterol metabolism. Further, eQTL in cluster 1 were mainly significant in the immune lineage and were enriched for genes contributing to cholesterol homeostasis, reflecting the central role of cholesterol metabolism in immune functions.^16^ Cluster 7, also mainly active in the immune lineage, was enriched for genes involved with, e.g., lipid transport. Lipid mediators are known to have an important role in lung fibrosis.^17^ The eQTL in cluster 2 showed opposing effects between the epithelial and immune lineages and were enriched for genes associated with highly lineage-specific functions, such as epithelial cell morphogenesis.

### Disease-specific eQTL are highly cell type-specific

To identify eQTL that are specific to healthy or affected individuals or that show a different direction or degree of effect in the two groups, we performed disease-state interaction eQTL (int-eQTL) mapping (**Methods**). Testing across 33 cell types with ≥5 ILD and ≥5 unaffected donors and a MAF ≥5% in each group, we detect 83,596 int-eQTL. Compared to the non-interaction-eQTL, there was substantially less lineage and cell type sharing of int-eQTL (**Fig. 5a, Supplementary Fig. 5**): for each gene, there was a median of 21 top int-eQTL (min=2, max=28), resulting in a total of 75,482 top int-eQTL. Compared to the top non-interaction sc-eQTL, int-eQTL were further from the TSS (mean absolute distance, sc-eQTL=43.1Mb, int-eQTL=52.9Mb, t-test *p*=2.22×10^−16^) and had larger effect sizes (mean absolute mashr posterior beta, sc-eQTL=0.10, int-eQTL=0.66, t-test *p*=2.22×10^−16^, **Fig. 5b**). Some disease int-eQTL were linked to overall expression differences between groups (**Fig. 5c**): 43% of int-eGenes were differentially expressed (adj. *p*<0.1) between ILD and unaffected samples in the particular cell type. Out of these genes, 50.8% were expressed at a higher level in ILD. However, 21% of int-eGenes were widely expressed (>30% of cells) in both groups in the particular cell type and did not exhibit notable differences in expression levels (|logFC|<0.2), indicating that these eGenes are equally expressed but are differentially affected by *cis*-regulatory loci. These include *DSP* with 3 top int-eQTL, including rs2003916, which showed differential effects between ILD and unaffected donors in four of the tested epithelial cell types (**Fig. 5d**).

**Fig. 5:**
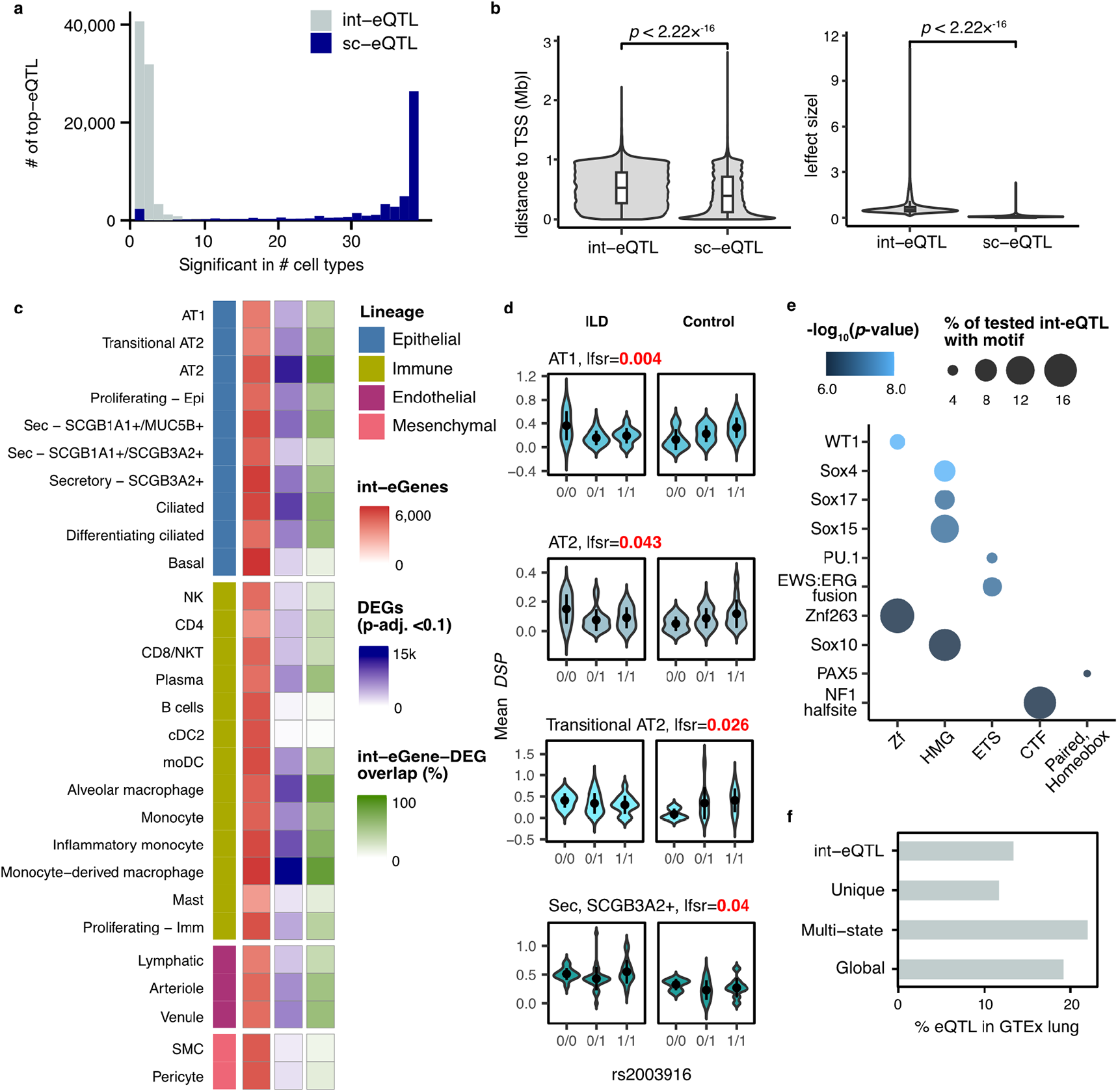
Disease-interaction eQTL converge in pathways relevant to lung fibrosis. **a**, Histogram of the cell type sharing of top int-eQTL and top non-interaction sc-eQTL. **b**, Comparison of absolute distances to the eGene TSS and absolute effect sizes of top sc-eQTL and int-eQTL. *t*-test *p*-values are indicated. **c**, Numbers of int-eGenes, DEGs between fibrotic and unaffected samples, and the proportion of their overlap for each cell type included in the int-eQTL analysis. **d**, An example of an int-eQTL for *DSP*. **e**, Top TF motifs enriched among int-eSNPs associated with eGenes that were equally expressed between ILD and unaffected donors but exhibited differences in eQTL effect sizes. TFs are grouped by family on the x-axis. **f**, Percentage of int-eQTL, sc-eQTL unique to a single cell type, multi-cell type sc-eQTL, and globally shared sc-eQTL that are also eQTL in GTEx lung (*p*<1×10^−6^).

To further interrogate the mechanisms underlying these int-eQTL, we analyzed the int-eQTL associated with eGenes expressed equally between ILD and unaffected donors for the enrichment of known transcription factor binding sites (TFBS, **Methods**). We identified 42 significantly enriched TF motifs (*q*<0.05), including WT1, several SOX, HOX, and PAX family members, ERG, and NF1 (**Fig. 5e, Supplementary Table 4**). Several of these have known importance in lung fibrosis. WT1 functions as a positive regulator of fibroblast proliferation, myofibroblast transformation, and ECM production.^18^ A number of SOX TFs are known to be upregulated in IPF and are associated with fibroblast activation.^19,20^ Out of the 37 genes encoding TFs disrupted by int-eQTL that were also tested for differential expression, 30 were DE between ILD and unaffected samples in at least one cell type. When examined across all cell types with significant DE, 43.0% of these genes were expressed at a higher level in ILD. The 7 that were equally expressed between cases and controls – including WT1, SOX10, PAX7, HOXA11, HOXD12, NKX6-1, and SCRT1 – could contribute to ILD pathogenesis through differences in protein levels or localization, differential binding to *cis*-regulatory elements, or chromatin level differences in addition to or instead of differential TF abundance.

We assessed the level at which sc-eQTL and int-eQTL are replicated in bulk analyses by overlapping the eQTL detected here with lung eQTL from GTEx.^2^ All classes of sc-eQTL as well as int-eQTL were enriched among GTEx lung eQTL (Fisher’s exact test, p<2.2×10^−16^). Out of the globally shared and multi-cell type top-eQTL, 19.1% and 21.9% were also eQTL in GTEx lung with a nominal p<1×10^−6^ (**Fig. 5f**). However, only 11.7% of sc-eQTL unique to a single cell type and 13.4% of int-eQTL were GTEx significant. This finding demonstrates the power of cell type and context-specific analyses in uncovering regulatory effects concealed by less granular approaches.

### Cell type-specific patterns of colocalization at GWAS loci

To connect the shared and cell type-specific regulatory variants to IPF risk, we compared our results to a recent IPF GWAS meta-analysis.^9^ All major classes of eQTL were enriched among loci implicated (nominal *p*<1×10^−6^, **Supplementary Table 5**) by the IPF GWAS meta-analysis (Fisher’s exact test, globally shared *p*<5.09×10^−64^, multi-cell type *p*<1.83×10^−98^, unique to a single cell type *p*=0.0525), while a null set of non-significant eQTL with a matched distribution of distances to TSS was not (*p*=1). GTEx bulk lung-eQTL were similarly highly enriched (*p*=2.22×10^−111^) among IPF GWAS loci. Disease-interaction eQTL, however, were not more enriched among IPF GWAS loci than a null set of non-significant eQTL.

In addition to the intersection analysis described above, we colocalized eQTL signals for 2,092 genes – including the target genes of the multi-state eQTL in **Fig. 4** and 103 GWAS implicated genes – with the IPF GWAS meta-analysis,^9^ the UK BioBank (UKBB) IPF GWAS,^21^ and an East Asian IPF GWAS^22^ (**Methods**). Here we identified 5 loci with evidence of colocalization (posterior probability for a single shared causal variant >0.6) between risk loci and eQTL in at least one cell type. These patterns largely overlapped between the IPF GWAS meta-analysis and UKBB (**Fig. 6, Supplementary Table 6**). Three of these loci were eQTL for genes that have previously been implicated in GWAS, *MUC5B, DSP*, and *KANSL1* according to the NHGRI EBI GWAS Catalog.^23^ The loci associated with *KANSL1* in both the GWAS and eQTL analysis was also associated with the expression of *KANSL1-AS1* across a number of cell types in our dataset. Additionally, we found an eQTL for the gene *JAML* was significantly colocalized with a locus from the GWAS analysis. This variant did not meet the criterion for genome-wide significance in the GWAS analysis, but was an eQTL across a number of myeloid lineage cell types (**Supplementary Fig. 10**). *MUC5B* was robustly expressed and colocalized with the IPF GWAS meta-analysis and the UKBB IPF GWAS in SCGB1A1+/MUC5B+ and SCGB3A2+ secretory cells, implicating these as the most likely cell types in which the risk variant functions (**Supplementary Fig. 7–9**). In contrast to the mostly European IPF GWAS meta-analysis and UKBB, *MUC5B* eQTL did not significantly colocalize with the East Asian IPF GWAS in any cell type, likely due to the low frequency of the risk allele in Asian populations.^24^ The pattern of population sharing was different for *DSP* eQTL, which was colocalized with the IPF GWAS meta-analysis in AT2, transitional AT2, and AT1 cells, and with UKBB and the East Asian IPF GWAS in AT2 cells (**Supplementary Fig. 11**). eQTL for *KANSL1* colocalized with the meta-analysis and UKBB in ciliated epithelial cells. Additionally, eQTL for *KANSL1-AS1* antisense RNA was widely colocalized with the meta-analysis and UKBB across epithelial, immune, and endothelial cell types. However, expression levels and eQTL effect sizes of *KANSL1* and *KANSL1-AS1* were highly correlated (**Supplementary Fig. 12–14**) and both genes are ubiquitous but lowly expressed across cell types, impeding an exact evaluation of the cell type specificity of these effects.

**Fig. 6:**
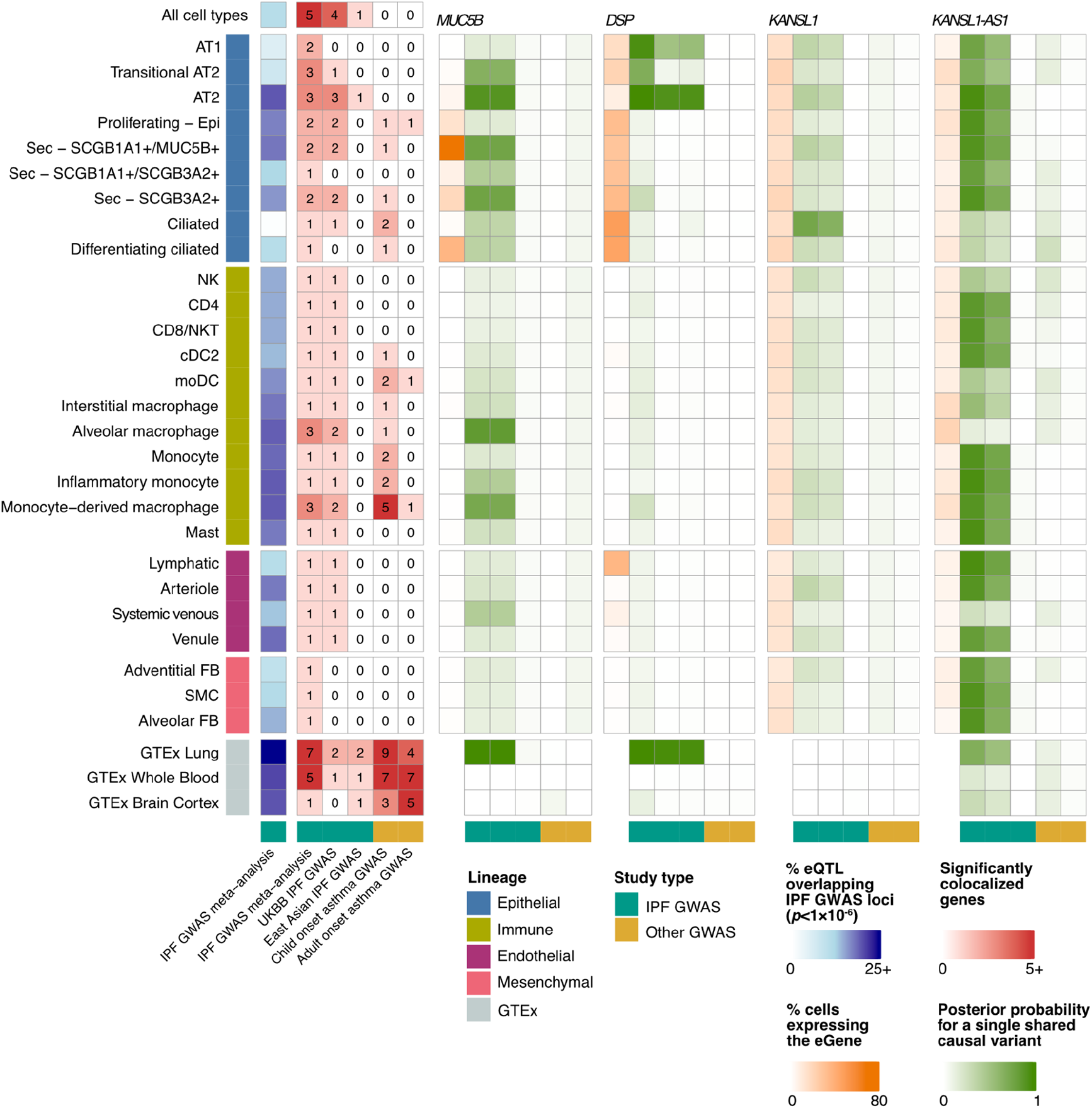
Cell type-eQTL colocalize with lung trait GWAS. Heatmaps presenting the numbers of eQTL that are nominally significant (*p*<1×10^−6^) in the IPF GWAS meta-analysis (blue), numbers of significant colocalizations between cell type and bulk-eQTL and three IPF GWASs, as well as child and adult-onset asthma GWAS (red), the proportion of cells expressing the gene (orange) and posterior probabilities for a single shared causal variant between the tested cell types and GWAS for selected top IPF associated genes (*MUC5B, DSP, KANSL1, KANSL1-AS1*, green) across 27 cell types with at least one colocalized gene.

When examining how these signals were colocalized in bulk eQTL analyses, we found colocalization patterns of *MUC5B* and *DSP* between GTEx lung and IPF GWAS reflect those of the cell type-level analysis (**Supplementary Fig. 6, Supplementary Table 6**): *MUC5B* was significantly colocalized with the IPF GWAS meta-analysis and UKBB, but not with the East Asian IPF GWAS. *DSP* was colocalized in all three IPF GWAS. *KANSL1*, however, did not colocalize between GTEx lung and any IPF GWAS. To assess to what extent the genetic and cell type-specific regulatory architecture of IPF risk may be shared with other lung diseases, we colocalized the cell type-eQTL signals with child and adult-onset asthma GWAS.^25^ The childhood-onset asthma colocalization revealed a regulatory architecture distinct from IPF, with a lack of colocalizations in epithelial cells and most of the significant colocalizations being specific to immune cells, particularly monocytes and monocyte-derived macrophages, which may shape some of the clinical and inflammatory features of asthma.^26,27^ These results highlight the broader utility of this dataset in the investigation of other lung traits and diseases.

## Discussion

Here, we present the first characterization of regulatory variants across major cell types in the human lung, employing scRNA-seq to identify eQTL at cell type resolution. In total, we characterized eQTL across 38 different cell types identifying *cis* eQTLs in over 6,000 genes. Building upon bulk eQTL studies, such as GTEx, which sought to characterize differences in gene regulatory architecture across tissues, we employed a multivariate adaptive shrinkage approach to robustly identify shared and specific eQTL across cell types.^2^ In addition to, the majority of eQTL which were shared across cell types, we identified thousands of eQTL that were limited to a subset or single cell type. These eQTL classes were enriched among chronic lung disease GWAS loci and differentially-expressed genes in PF lungs, suggesting that context-specific gene regulatory mechanisms are important yet to-date largely unrecognized contributors to the mechanisms underlying chronic lung diseases.

Highlighting the power of this approach, we demonstrate that many of the eQTL identified here were not eQTL in bulk data from primary lung tissue (**Fig. 5f**). This was particularly true of eQTL limited to a single cell type (11.7% significant in bulk) and disease-interaction eQTL, which are far less likely to be shared across cell types (13.4% significant in bulk). Both of these classes of eQTL tend to be further away from the TSS than global and multi-state eQTL suggesting these loci may be disrupting enhancers rather than promoters (**Fig. 4a, Fig. 5b**). This observation would be consistent with the cell type specificity of these eQTL and would distinguish them from eQTL identified in bulk studies, which are strongly enriched for disrupting promoter regions. Indeed, some work suggests that common eQTL (enriched near promoters) are less likely to have functional relevance^1,28,29^. In addition to being more distal from the TSS, these eQTL also tend to have larger effect sizes (**Fig. 4a, Fig. 5b**). At present, it is uncertain whether the difference in effect size is due to statistical power to identify these associations or if cell type specific eQTL inherently exhibit larger effect sizes. Since this class of eQTL is the least likely to benefit from the mashr^12^ approach, it seems plausible that we only have statistical power to identify those with large effects. If this is the case, future single cell eQTL studies with increased sample number and cell type representation from rare cell populations are likely to identify a significant number of additional cell type and context specific eQTL.

Over the past ten years, there has been an increased appreciation for the degree to which eQTL may be context specific, starting first with tissue type, then to functional/environmental context, and finally to cell type^30–37^. The results of this study suggest single cell eQTL studies have the power to more robustly elucidate this context specificity, and that they will better recover eQTL associated with disease states or environmental perturbations as these effects are less likely to be shared across cell type within a tissue.

In addition to a general characterization of eQTL in the lung, this study is uniquely positioned to explore the interplay between genetic variation and the molecular underpinnings of chronic lung diseases including PF. Focusing first on the known risk loci identified in various GWAS studies, we found eQTL to be enriched among GWAS risk loci regardless of class (**Fig. 6**). These enrichments were similar to those found in bulk eQTL analysis from the human lung (**Fig. 6**), however using the cell type level associations, we are able to partition the function of these risk variants into discrete cell types. Indeed, we find risk variants are most likely to be eQTLs in AT2 cells, followed by a number of cells from the myeloid lineage – including both resident and recruited macrophages (**Fig. 6, Supplementary Table 5**). Using a more formal colocalization analysis we found four GWAS loci with strong support for a shared causal variant with an eQTL (compared to seven colocalizations in the bulk eQTL data), for which we were able to identify the likely cell type in which these risk variants are acting (**Fig. 6**). Our findings align with recent insights into the cellular and regulatory drivers of ILD. Epithelial cell types are suggested to have a central role in driving alveolar remodeling in IPF.^38^ Indeed, in a GWAS colocalization analysis, we find that the top IPF risk variants flanking *MUC5B* and *DSP* regulate the expression levels of their targets in specific epithelial cell types.

In addition to assessing the effect of known risk loci on gene expression traits, we were also able to more directly examine how genetic variation may alter key regulatory processes involved in disease. Turning back to the disease-interaction eQTL analysis, enabled by the collection of a cohort composed of both diseased and unaffected individuals, we were able to assess how these context specific eQTL may further drive disease processes. Roughly half of the interaction eQTLs are driven by differences in overall mean expression between the diseased and control samples. In the case of disease emergent expression difference (expression increases in diseased sample), loci which further upregulate gene expression may propagate additional molecular dysfunction. Focusing on the set of interaction eQTL with similar mean expression across disease and control samples, we found the loci to be enriched for TFBS that are associated with key biological processes related to ILD. For example, we found enrichment for WT1^18^ and SOX family members^19,20^, which previous experimental evidence connects to fibroblast activation and proliferation in the lung. The eQTL which disrupt key binding sites likely further propagate the molecular dysregulation observed in ILD by modulating binding efficiency of TFs and altering expression of their direct and downstream target genes. Of note, interaction eQTL were not enriched for overlaps with risk variants, as anticipated based on the presumed requirement for disease-associated contextual cues for these variants to manifest their effects.We postulate these context-specific eQTL may play a role in disease progression rather than initiation. Again, these results highlight the importance of identifying context specific eQTL which are best captured using single cell approaches.

Taken together, our study demonstrates the powerful application of single-cell genomics to study genetic regulation of gene expression in complex, solid, primary human tissues. Integrating scRNA-seq data from control and diseased lung samples with genetic data provides new insights into the cell-type specific function of risk variants for interstitial lung disease, and highlights a new class of regulatory variants (interaction eQTL) that contribute to disease pathobiology. Future work combining single-cell ‘omic assays, healthy and disease samples, and context-specific analysis methods will be important to understanding the interplay of dysfunctional genetic regulation and cellular contexts in complex human disease.

## Acknowledgments

We thank Tennessee Donor Services and the Donor Network of Arizona and the patients and families who donated tissue samples to make these studies possible. This study was supported by NHLBI R01HL145372 and DOD W81XWH1910416, to N.E.B. and J.A.K., P01HL092870 to T.S.B., K08HL136888 to C.M.S., NHGRI R01HG011886 to N.E.B, D.J.M, and J.A.K., the Doris Duke Charitable Foundation to J.A.K. and N.E.B., and NHMRC GNT1195595 and GNT1162829 to D.J.M. The Vanderbilt Flow Cytometry Shared Resource is supported by the Vanderbilt Ingram Cancer Center (P30 CA068485) and the Vanderbilt Digestive Disease Research Center (DK058404).

## Methods

### Subjects, samples, and tissue processing

The data presented here include 82 previously published^39^ samples from 56 individuals and 83 previously unpublished samples (**Supplementary Table 1**). In addition to previously published data, lung tissue samples were collected from 60 individuals, including 33 ILD cases and 27 nonfibrotic controls, and processed as previously described by Habermann et al.^8^ Briefly, ILD tissue samples were obtained from lungs removed at the time of lung transplantation at either Vanderbilt University Medical Center (VUMC) or the National Thoracic Institute (NTI). Control tissue samples were obtained from lungs declined for organ donation either at the Donor Network of Arizona (DNA) or VUMC. Tissue sections were taken from multiple peripheral (within ∼2 cm of the pleural surface) regions in each lung. For ILD lungs, representatively diseased areas were selected on the basis of preoperative chest CT, while for control lungs, the most normal-appearing region was identified by gross inspection and selected for biopsy. For ILD lungs, diagnoses were determined according to the American Thoracic Society/European Respiratory Society consensus criteria^40^. Studies were approved by the local Institutional Review Boards (Vanderbilt IRB nos. 060165 and 171657 and Western IRB no. 20181836).

Tissue samples were digested in either collagenase I/dispase II (1 μg/ml) or Miltenyi Multi-Tissue Dissociation Kit using a gentleMACS Octo Dissociator (Miltenyi Inc.). Tissue lysates were serially filtered through sterile gauze, 100- and 40-μm sterile filters (Fischer). The resulting suspensions then underwent cell sorting using serial columns (Miltenyi Microbeads, CD235a and CD45) or fluorescence-activated cell-sorting (FACS) at VUMC or FACS at the Translational Genomics Research Institute (TGen). CD45^−^ and C45^+^ populations were mixed 2:1 in samples processed at VUMC and used to generate the scRNA-seq libraries. At TGen, calcein acetoxymethyl was used to stain live cells, and 10,000 to 15,000 live cells were sorted directly into the 10x reaction buffer and transferred to the 10x 5′ chip A (10x Genomics).

### scRNA-seq library preparation and next-generation sequencing

scRNA-seq libraries were generated using the 10x Chromium platform 5′ library preparation kits (10x Genomics) according to the manufacturer’s recommendations and targeting 5000 to 10,000 cells per sample. From 12 donors, multiple tissue samples were processed and libraries were generated from separate biopsies taken from the same lung to account for regional heterogeneity (**Supplementary Table 1**). Next-generation sequencing was carried out on an Illumina NovaSeq 6000 or HiSeq 4000. The resulting sequence data were filtered to retain reads with a read quality >3, and CellRanger Count v3.0.2 (10x Genomics) was used to align reads onto the GRCh38 reference genome.

### Data integration, clustering, and cell type annotation

scRNA-seq data were processed and analyzed using Seurat v4.^10^ CellRanger Count outputs were imported to create a Seurat object for each sample. The sample-specific objects were merged and the proportions of reads arising from mitochondrial genes were calculated for each sample. The merged object was filtered to retain samples with more than 1,000 identified features or less than 25% of mitochondrial reads.

Samples sequenced across 24 batches were integrated using reciprocal principal component analysis (rPCA) as follows: The merged object was split by flowcell and the count data in each batch-specific object was normalized. Variable features were identified for each object, and integration features across objects were selected with SelectIntegrationFeatures(). Data in each batch-specific object was scaled and underwent PCA dimensionality reduction using 2,000 variable features. rPCA integration was carried out using 3,000 integration anchors and four reference batches (6, 12, 18, 24). PCA dimensionality reduction on the integrated data was performed using 3,000 variable features. To determine the optimal number of PCs to identify neighbors and to construct the UMAP, we determined the difference between the variation explained by each PC and the subsequent PC and identified the last point where the percentage change was more than 0.1%. A Shared Nearest Neighbor graph was constructed with k=20, and clusters of cells were identified using the modularity optimization-based clustering algorithm^41^ implemented in Seurat v4.

The resulting clusters were divided into four major cell subgroups based on marker gene expression: *PTPRC*^+^ for immune cells, *EPCAM*^*+*^ for epithelial cells, *PECAM1*^*+*^*/PTPRC*^*−*^ for endothelial cells, and *PTPRC*^*−*^*/EPCAM*^*−*^*/PECAM1*^*−*^ for mesenchymal cells. Each subgroup-specific object underwent the same dimensionality reduction and clustering approach as described above. Clusters containing doublets were removed by identifying clusters of cells that expressed markers specific to multiple cell lineages. After doublet removal and reclustering, the subgroup-specific objects were further annotated for specific cell types based on known marker genes (**Supplementary Table 2**).

### Differential gene expression and cell type proportion testing

We tested for differences in cell type abundances between groups using the propeller.anova() function of R/speckle.^42^ For differential gene expression, we used the R/presto implementation of the Wilcoxon rank sum test (wilcoxauc).^43^ For overlapping with disease-interaction eGenes, a relaxed significance threshold of adj. *p* <0.1 was employed.

### Low-pass Whole Genome Sequencing, genotyping, and imputation

Flash-frozen tissue in DNA/RNA Shield was homogenized via a bullet blender. Genomic DNA was extracted using Zymo Quick DNA/RNA microprep plus kit. Library preparation and low-pass Whole Genome Sequencing were carried out at TGen or by Gencove (**Supplementary Table 7**), Inc. At TGen, libraries were prepared using PCR-free Watchmaker kits (Watchmaker Genomics) with 200 ng input. Genomes were sequenced on NovaSeq at low coverage (typically 0.4–1x). The resulting sequence data were processed and imputed using Gencove’s imputation platform.

### Pseudo-bulk cell type eQTL mapping

For eQTL mapping, cells with >20% of reads mapping to the mitochondrial genes were removed (466,989 cells remain). Mapping was only performed on cell types with at least 40 donors with at least 5 cells of that cell type (38 cell types meet these criteria). Donor VUILD65 was removed due to inconsistencies in metadata suggesting mislabeling. Mitochondrial genes, genes encoding ribosomal proteins (downloaded from HGNC: https://www.genenames.org/cgi-bin/genegroup/download?id=1054&type=branch), genes expressed in fewer than 10% of cells in the study, and genes with a mean count across all cells < 0.1 were excluded, resulting in 6,995 genes for eQTL mapping.

Pseudobulk cis-eQTL mapping was performed following guidelines from Cuomo, Alvari, Azodi et al. 2021^11^. For each cell type, raw counts were normalized and log_2_ transformed using scran^44^ and mean aggregated to get a single value for each gene for each donor for each cell type. Donors with <5 cells for a cell type were excluded from eQTL mapping for that cell type and only cell types with at least 40 donors matching this criteria were included (max donors=113). Biallelic, autosomal SNPs were filtered to include SNPs with a minor allele frequency > 5%, Hardy-Weinberg equilibrium *p*>1×10^−6^, and further pruned to remove highly correlated SNPs (--indep-pairwise 250 50 0.9) using plink2,^45^ resulting in ∼1.9 million SNPs. We tested for associations for SNPs within 1Gb up and downstream of the gene body.

Linear mixed models were used to map *cis-*eQTL using the LIMIX_qtl framework (https://github.com/single-cell-genetics/LIMIX_qtl).^46^ Expression levels for each gene were quantile-normalized to fit a normal distribution (--gaussianize_method). To control for unwanted technical effects, the first 20 cell type expression principal components (PCs) were regressed out before model fitting (--regress_covariates). To account for variance due to population structure, we included the identity-by-descent (IBD) relationship matrix generated by applying plink2 --make-rel on the filtered SNP data as a random effect. To account for differences in cell type abundance across donors, we included the number of cells aggregated (1/nCells) as a second random effect, using the random effect weighting approach described by Cuomo, Alvari, Azodi, et al^11^. Random effects were marginalized from the model using the low-rank optimization method (--low_rank_random_effect) described by Cuomo et al.^47^

### Joint cell type eQTL analysis

Joint analysis of the LIMIX estimated effect sizes and their corresponding standard errors across all 38 cell types was performed using multivariate adaptive shrinkage in R (mashr v0.2 ^12^) following the approach outlined in the ‘eQTL analysis outline’ vignette from the authors (https://stephenslab.github.io/mashr/articles/eQTL_outline.html). In this approach, a weighted combination of learned and canonical covariance matrices that describe patterns of eQTL sparsity and sharing across cell types is used as a prior for generating adjusted summary statistics. The data-driven covariance matrices were estimated from a subset of strong associations with a local false sign rate < 0.1 in at least one cell type (n=487), calculated using adaptive shrinkage in R (ashr v2.2^48^). The default canonical covariance matrices were used, representing equal effect sharing across cell types, the top 5 PCs from the strong associations, and extreme deconvolution matrices obtained from those PCs. The model was fit to a random subset of 10,000 SNP-gene associations and then applied to all associations tested.

### Assessing significance, sharing, and eQTL classification

The local false sign rate (lfsr) calculated by mashr was used to assess significance. To further reduce the impact of differential power on assessing sharing of eQTL across cell types, if an eQTL was significant in one cell type (lfsr ≤0.05), then it would be considered significant in other cell types at a less stringent threshold (lfsr ≤0.1). An eQTL was considered shared in a pairwise comparison between two cell types if the eQTL was significant in both cell types and the estimated effect size was within a factor of 0.5. An eQTL was classified as global if it was significant in at least 36 of the 38 cell types (31 of 33 cell types for interaction eQTL). This two-cell-type buffer was included to reduce the impact of low-powered cell types on our categorization. eQTL that were significant in only one cell type were classified as unique, and eQTL significant in 2-36 cell types (2-31 for interaction eQTL) were considered multi-cell type eQTL.

To simplify plotting of top-eQTL, a pruning step is included, where for each gene, if there is a single top-eQTL, that eQTL is retained. If there are two top-eQTL, the Euclidean distance between the centered absolute values of the estimated effect sizes across cell types for the two eQTL are compared. If the distance is greater than the set threshold (dist=0.2), both are retained. If the distance is less than the threshold then the one that is significant in more cell types is retained. Finally, if there are more than three top-eQTL, the pairwise Euclidean distance between the centered absolute values of the estimated effect sizes for each pair of top-eQTL is calculated. If all pairwise distances are above the threshold, all are retained. Otherwise, hierarchical clustering is performed and the tree is cut using cutree at k between 2 and 5 that maximizes the Silhouette width. For each cluster, the top-eQTL that is significant in most cell types is retained.

### Disease-interaction cell type-eQTL mapping

To test for disease-interaction eQTL effects, cell types were required to have at least 10 control and 10 ILD donors with at least 5 cells of that cell type, resulting in KRT5-/KRT17+, pDC, cDC1, alveolar FB, and mesothelial cell types being excluded from the interaction eQTL analysis. SNPs were further filtered to remove those with a MAF < 5% in either the control or ILD donor populations (1.77 million SNPs remained). Interaction effects were tested using the run_interaction_QTL_analysis from LIMIX_qtl. Random effects were handled as described above for the eQTL mapping analysis. In the interaction term with SNP effect, we included the binary disease status (ILD vs. unaffected). Fix effects (to 20 PCs) were included, but not regressed out before modeling, as disease status was strongly correlated with some PCs. The results from this analysis were processed using mashr, with significance calling, as described above for the eQTL analysis. For each cell type, we further pruned the int-eQTL to retain associations where the observed eSNP MAF for ILD and unaffected donors for the given cell type was >0.05.

### Colocalization with GWAS and GTEx

Colocalization analysis was carried out between the cell type eQTL, GTEx lung eQTL, and three IPF genome-wide association studies (GWAS). UK Biobank (UKBB)^21^ and East Asian^22^ IPF GWAS summary statistics were downloaded from GWAS Catalog.^23^ The discovery samples of these studies comprised 1,369 IPF cases, 14,103 COPD cases and 435,866 controls, and 1,046 East Asian ancestry cases and 176,974 controls, respectively. Summary statistics from an IPF GWAS meta-analysis^9^ leveraging data from three studies^49–51^ were downloaded after gaining access through submitting a request (https://github.com/genomicsITER/PFgenetics). The meta-analysis comprised 2,668 European ancestry IPF cases and 8,591 controls.

Additionally, GWAS on adult and child-onset asthma^25^ (26,582 adult European ancestry cases, 13,962 child cases, 300,671 controls) were downloaded from GWAS Catalog and included for comparison. For comparative analyses with bulk-eQTL, GTEx lung, whole blood, and brain cortex eQTL summary statistics were downloaded from the GTEx Google Cloud bucket (https://console.cloud.google.com/storage/browser/gtex-resources).

Bayesian colocalization analysis was performed using R/coloc v5.^52^ For the pseudo-bulk cell type-eQTL, mashr posteriors were used in place of nominal eQTL summary statistics. 2,092 genes, including the multi-cell type eQTL presented in **Fig. 4** and 103 IPF GWAS variant flanking genes, were selected for the colocalization analysis, and for each gene, colocalization testing was carried out between datasets that shared ≥100 variable (MAF >0, <1) SNPs. Significantly colocalized loci were selected based on the posterior probability for a single shared causal variant ≥0.6.

### Enrichment testing

We tested for the enrichment of the clusters of eQTL in Fig. 4 among Gene Ontology terms using R/TopGO v2.46.0.^53^ All genes included in the eQTL analysis were used as a background set. A *p*-value threshold of 0.01 was used to select significant terms. For the GO overrepresentation testing of the GTEx colocalizing genes, we used R/gprofiler2^54^ and *p*<0.05. Both are implementations of Fisher’s test.

We used Fisher’s exact test to test for the enrichment of the various classes of sc-eQTL (all eQTL, globally shared, multi-state, unique to a single cell type, k1-k7 in **Fig. 4**) among IPF GWAS risk variants. From the 1,617,891 SNPs tested for in the eQTL analysis and included in the IPF GWAS meta-analysis, a set of 473 GWAS variants was selected with a relaxed genome-wide nominal p-value threshold of 1×10^−6^. A null distribution of non-significant eQTL was generated using the default rejection method of R/nullranges^55^ to match the observed distribution of absolute distances to TSS among the significant eQTL.

To test whether the various classes of regulatory variants detected in the sc-eQTL analyses disrupt the binding of known transcription factors (TFs), we used HOMER^56^ to analyze eQTL positions for the enrichment of TF binding site motifs. findMotifsGenome.pl with the default region size of 200bp was used to detect enriched motifs. In each analysis, a null set of non-significant eQTL with a matched distribution of distances to the transcription start site (TSS) was used as a background. In the TFBS enrichment analysis of the int-eQTL, the non-interaction sc-eQTL were used as a background set. A *q*-value threshold of 0.05 was used to select significant motifs.

## Data and code availability

Raw and processed 10x Genomics data, Seurat objects, mean-aggregated expression matrices, and genome-wide LIMIX and mashr eQTL statistics can be found on GEO with the accession number GSE227136. Genotype data are being made available on dbGaP. Custom scripts to reproduce the result presented here are available on GitHub at https://github.com/tgen/banovichlab/tree/master/ILD_eQTL.

## References

1. Umans, B. D., Battle, A. & Gilad, Y. Where Are the Disease-Associated eQTLs? Trends Genet. 37, 109–124 (2021).

2. Consortium, T. G. & The GTEx Consortium. The GTEx Consortium atlas of genetic regulatory effects across human tissues. Science vol. 369 1318–1330 Preprint at https://doi.org/10.1126/science.aaz1776 (2020).

3. Lea, A. J., Peng, J. & Ayroles, J. F. Diverse environmental perturbations reveal the evolution and context-dependency of genetic effects on gene expression levels. Genome Res. (2022) doi:10.1101/gr.276430.121.

4. GTEx Consortium. Genetic effects on gene expression across human tissues. Nature 550, 204 (2017).

5. Lederer, D. J. & Martinez, F. J. Idiopathic Pulmonary Fibrosis. New England Journal of Medicine vol. 378 1811–1823 Preprint at https://doi.org/10.1056/nejmra1705751 (2018).

6. Ley, B., Collard, H. R. & King, T. E., Jr. Clinical course and prediction of survival in idiopathic pulmonary fibrosis. Am. J. Respir. Crit. Care Med. 183, 431–440 (2011).

7. Moss, B. J., Ryter, S. W. & Rosas, I. O. Pathogenic Mechanisms Underlying Idiopathic Pulmonary Fibrosis. Annu. Rev. Pathol. 17, 515–546 (2022).

8. Habermann, A. C. et al. Single-cell RNA sequencing reveals profibrotic roles of distinct epithelial and mesenchymal lineages in pulmonary fibrosis. Sci Adv 6, eaba1972 (2020).

9. Allen, R. J. et al. Genome-Wide Association Study of Susceptibility to Idiopathic Pulmonary Fibrosis. Am. J. Respir. Crit. Care Med. 201, 564–574 (2020).

10. Hao, Y. et al. Integrated analysis of multimodal single-cell data. Cell 184, 3573–3587.e29 (2021).

11. Cuomo, A. S. E. et al. Optimizing expression quantitative trait locus mapping workflows for single-cell studies. Genome Biol. 22, 188 (2021).

12. Urbut, S. M., Wang, G., Carbonetto, P. & Stephens, M. Flexible statistical methods for estimating and testing effects in genomic studies with multiple conditions. bioRxiv 096552 (2018) doi:10.1101/096552.

13. Dimas, A. S. et al. Common Regulatory Variation Impacts Gene Expression in a Cell Type– Dependent Manner. Science vol. 325 1246–1250 Preprint at https://doi.org/10.1126/science.1174148 (2009).

14. Mu, Z. et al. The impact of cell type and context-dependent regulatory variants on human immune traits. Genome Biol. 22, 122 (2021).

15. Popmihajlov, Z. et al. CC-90001, a c-Jun N-terminal kinase (JNK) inhibitor, in patients with pulmonary fibrosis: design of a phase 2, randomised, placebo-controlled trial. BMJ Open Respir Res 9, (2022).

16. Aguilar-Ballester, M., Herrero-Cervera, A., Vinué, Á., Martínez-Hervás, S. & González-Navarro, H. Impact of Cholesterol Metabolism in Immune Cell Function and Atherosclerosis. Nutrients 12, (2020).

17. Suryadevara, V., Ramchandran, R., Kamp, D. W. & Natarajan, V. Lipid Mediators Regulate Pulmonary Fibrosis: Potential Mechanisms and Signaling Pathways. Int. J. Mol. Sci. 21, (2020).

18. Sontake, V. et al. Wilms’ tumor 1 drives fibroproliferation and myofibroblast transformation in severe fibrotic lung disease. JCI Insight 3, (2018).

19. Gajjala, P. R. et al. Dysregulated overexpression of Sox9 induces fibroblast activation in pulmonary fibrosis. JCI Insight 6, (2021).

20. Zhou, J. et al. microRNA-186 in extracellular vesicles from bone marrow mesenchymal stem cells alleviates idiopathic pulmonary fibrosis via interaction with SOX4 and DKK1. Stem Cell Res. Ther. 12, 96 (2021).

21. Duckworth, A. et al. Telomere length and risk of idiopathic pulmonary fibrosis and chronic obstructive pulmonary disease: a mendelian randomisation study. Lancet Respir Med 9, 285–294 (2021).

22. Sakaue, S. et al. A cross-population atlas of genetic associations for 220 human phenotypes. Nat. Genet. 53, 1415–1424 (2021).

23. Buniello, A. et al. The NHGRI-EBI GWAS Catalog of published genome-wide association studies, targeted arrays and summary statistics 2019. Nucleic Acids Res. 47, D1005– D1012 (2019).

24. Peljto, A. L. et al. The MUC5B promoter polymorphism is associated with idiopathic pulmonary fibrosis in a Mexican cohort but is rare among Asian ancestries. Chest 147, 460–464 (2015).

25. Ferreira, M. A. R. et al. Genetic Architectures of Childhood- and Adult-Onset Asthma Are Partly Distinct. Am. J. Hum. Genet. 104, 665–684 (2019).

26. van der Veen, T. A., de Groot, L. E. S. & Melgert, B. N. The different faces of the macrophage in asthma. Curr. Opin. Pulm. Med. 26, 62–68 (2020).

27. Niessen, N. M. et al. Neutrophilic asthma features increased airway classical monocytes. Clin. Exp. Allergy 51, 305–317 (2021).

28. Glassberg, E. C., Gao, Z., Harpak, A., Lan, X. & Pritchard, J. K. Evidence for Weak Selective Constraint on Human Gene Expression. Genetics 211, 757–772 (2019).

29. Mostafavi, H., Spence, J. P., Naqvi, S. & Pritchard, J. K. Limited overlap of eQTLs and GWAS hits due to systematic differences in discovery. bioRxiv 2022.05.07.491045 (2022) doi:10.1101/2022.05.07.491045.

30. Lonsdale, J. et al. The Genotype-Tissue Expression (GTEx) project. Nat. Genet. 45, 580– 585 (2013).

31. Strober, B. J. et al. Dynamic genetic regulation of gene expression during cellular differentiation. Science 364, 1287–1290 (2019).

32. Banovich, N. E. et al. Impact of regulatory variation across human iPSCs and differentiated cells. Genome Res. 28, 122–131 (2018).

33. Ward, M. C., Banovich, N. E., Sarkar, A., Stephens, M. & Gilad, Y. Dynamic effects of genetic variation on gene expression revealed following hypoxic stress in cardiomyocytes. Elife 10, (2021).

34. Resztak, J. A. et al. Genetic control of the dynamic transcriptional response to immune stimuli and glucocorticoids at single cell resolution. bioRxiv (2022) doi:10.1101/2021.09.30.462672.

35. Yazar, S. et al. Single-cell eQTL mapping identifies cell type-specific genetic control of autoimmune disease. Science 376, eabf3041 (2022).

36. Bryois, J. et al. Cell-type-specific cis-eQTLs in eight human brain cell types identify novel risk genes for psychiatric and neurological disorders. Nat. Neurosci. 25, 1104–1112 (2022).

37. Nathan, A. et al. Single-cell eQTL models reveal dynamic T cell state dependence of disease loci. Nature 606, 120–128 (2022).

38. Chakraborty, A., Mastalerz, M., Ansari, M., Schiller, H. B. & Staab-Weijnitz, C. A. Emerging Roles of Airway Epithelial Cells in Idiopathic Pulmonary Fibrosis. Cells 11, (2022).

39. Bui, L. T. et al. Chronic lung diseases are associated with gene expression programs favoring SARS-CoV-2 entry and severity. Nat. Commun. 12, 4314 (2021).

40. Travis, W. D. et al. An official American Thoracic Society/European Respiratory Society statement: Update of the international multidisciplinary classification of the idiopathic interstitial pneumonias. Am. J. Respir. Crit. Care Med. 188, 733–748 (2013).

41. Waltman, L. & van Eck, N. J. A smart local moving algorithm for large-scale modularity-based community detection. The European Physical Journal B vol. 86 Preprint at https://doi.org/10.1140/epjb/e2013-40829-0 (2013).

42. Phipson, B. et al. propeller: testing for differences in cell type proportions in single cell data. Bioinformatics 38, 4720–4726 (2022).

43. Korsunsky, I., Nathan, A., Millard, N. & Raychaudhuri, S. Presto scales Wilcoxon and auROC analyses to millions of observations. bioRxiv (2019) doi:10.1101/653253.

44. Lun, A. T. L., McCarthy, D. J. & Marioni, J. C. A step-by-step workflow for low-level analysis of single-cell RNA-seq data with Bioconductor. F1000Res. 5, 2122 (2016).

45. Purcell, S. et al. PLINK: a tool set for whole-genome association and population-based linkage analyses. Am. J. Hum. Genet. 81, 559–575 (2007).

46. Lippert, C., Casale, F. P., Rakitsch, B. & Stegle, O. LIMIX: genetic analysis of multiple traits. Preprint at https://doi.org/10.1101/003905.

47. Cuomo, A. S. E. et al. CellRegMap: A statistical framework for mapping context-specific regulatory variants using scRNA-seq. Preprint at https://doi.org/10.1101/2021.09.01.458524.

48. Stephens, M. False discovery rates: a new deal. Biostatistics 18, 275–294 (2017).

49. Peljto, A. L. et al. Association between the MUC5B promoter polymorphism and survival in patients with idiopathic pulmonary fibrosis. JAMA 309, 2232–2239 (2013).

50. Fingerlin, T. E. et al. Genome-wide association study identifies multiple susceptibility loci for pulmonary fibrosis. Nat. Genet. 45, 613–620 (2013).

51. Noth, I. et al. Genetic variants associated with idiopathic pulmonary fibrosis susceptibility and mortality: a genome-wide association study. Lancet Respir Med 1, 309–317 (2013).

52. Wallace, C. A more accurate method for colocalisation analysis allowing for multiple causal variants. PLoS Genet. 17, e1009440 (2021).

53. Adrian Alexa, J. R. topGO. (Bioconductor, 2017). doi:10.18129/B9.BIOC.TOPGO.

54. Kolberg, L., Raudvere, U., Kuzmin, I., Vilo, J. & Peterson, H. gprofiler2 --an R package for gene list functional enrichment analysis and namespace conversion toolset g:Profiler. F1000Res. 9, (2020).

55. Davis, E. S. et al. matchRanges: Generating null hypothesis genomic ranges via covariate-matched sampling. Preprint at https://doi.org/10.1101/2022.08.05.502985.

56. Heinz, S. et al. Simple combinations of lineage-determining transcription factors prime cisregulatory elements required for macrophage and B cell identities. Mol. Cell 38, 576–589 (2010).

