## Supplementary Information for "Cell type-specific and disease-associated eQTL in the human lung"

Supplementary Figures S1–S14

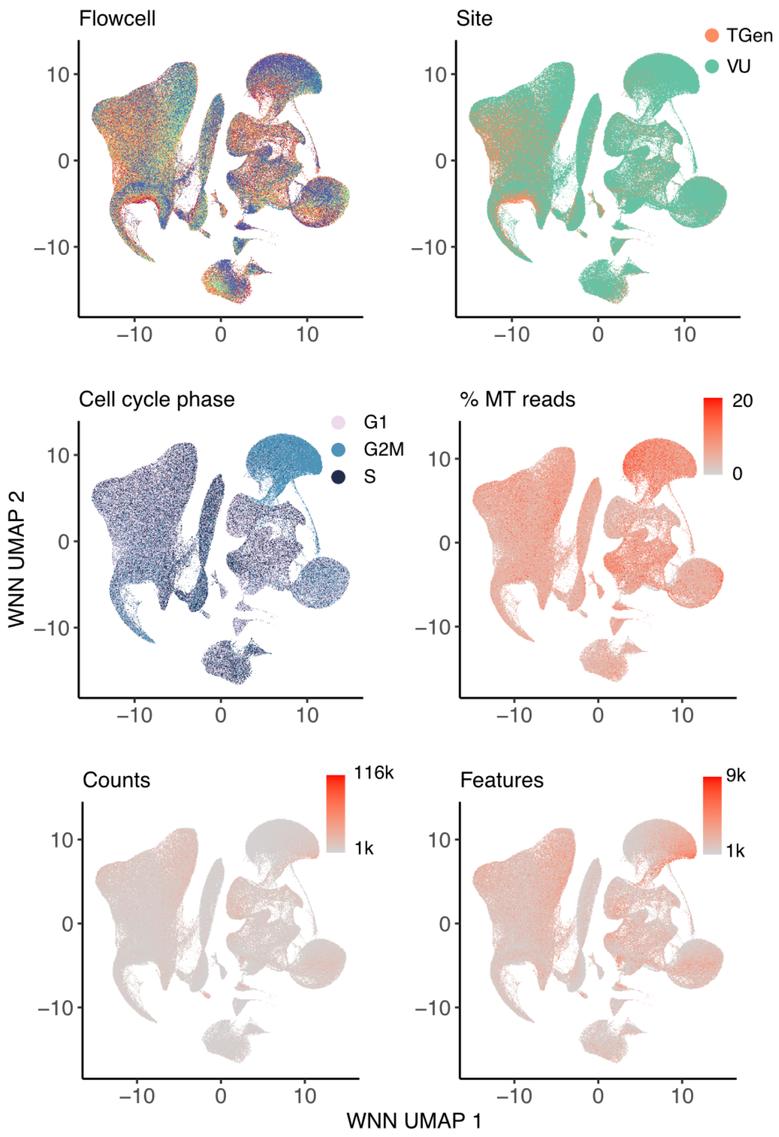


**Fig. S1:** UMAP dimensionality reductions of cells included in the pseudobulking and eQTL mapping, pseudocolored by flowcell, processing site (TGen or Vanderbilt), cell cycle phase, proportions of mitochondrial reads, number of read counts, and number of features.

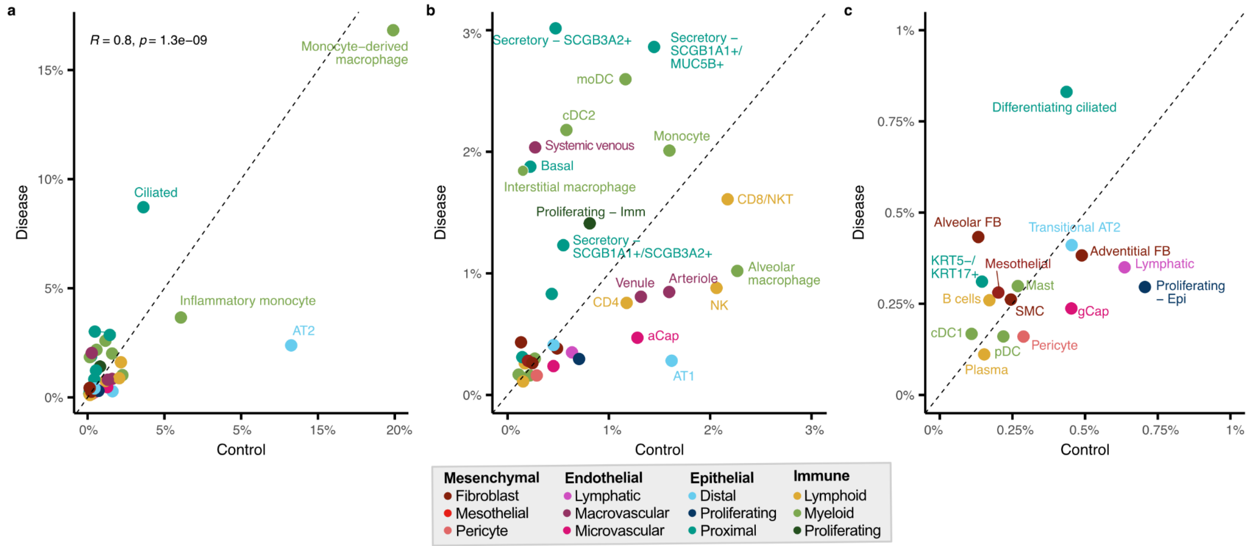


**Fig. S2:** Comparison of the median cell type abundance between control (x-axis) and diseased (y-axis) individuals for cell types included in eQTL mapping (# cell types = 38). The dashed line represents equal abundance in control and diseased samples. Cell types are colored by sub-lineage. **a,** All cell types shown with cell types with >3% abundance labeled. Pearson's correlation shown. **b,** Includes cell types with <3% abundance, with cell types with >1% abundance labeled. **c,** Includes and labels all cell types with <1% abundance.

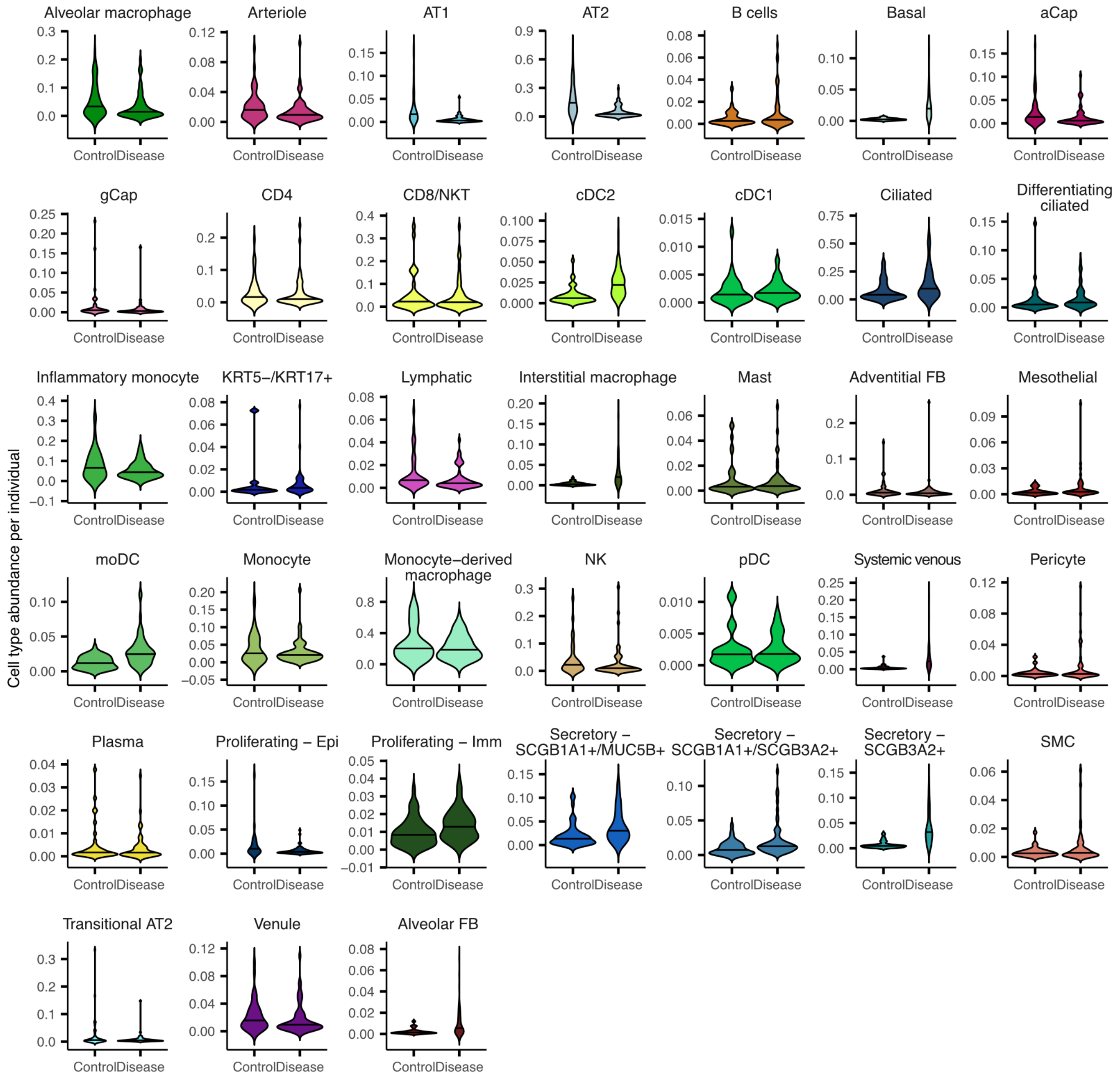


**Fig. S3:** The distribution of cell type abundance per individual grouped by disease status (control vs. diseased). The bar represents the median abundance.

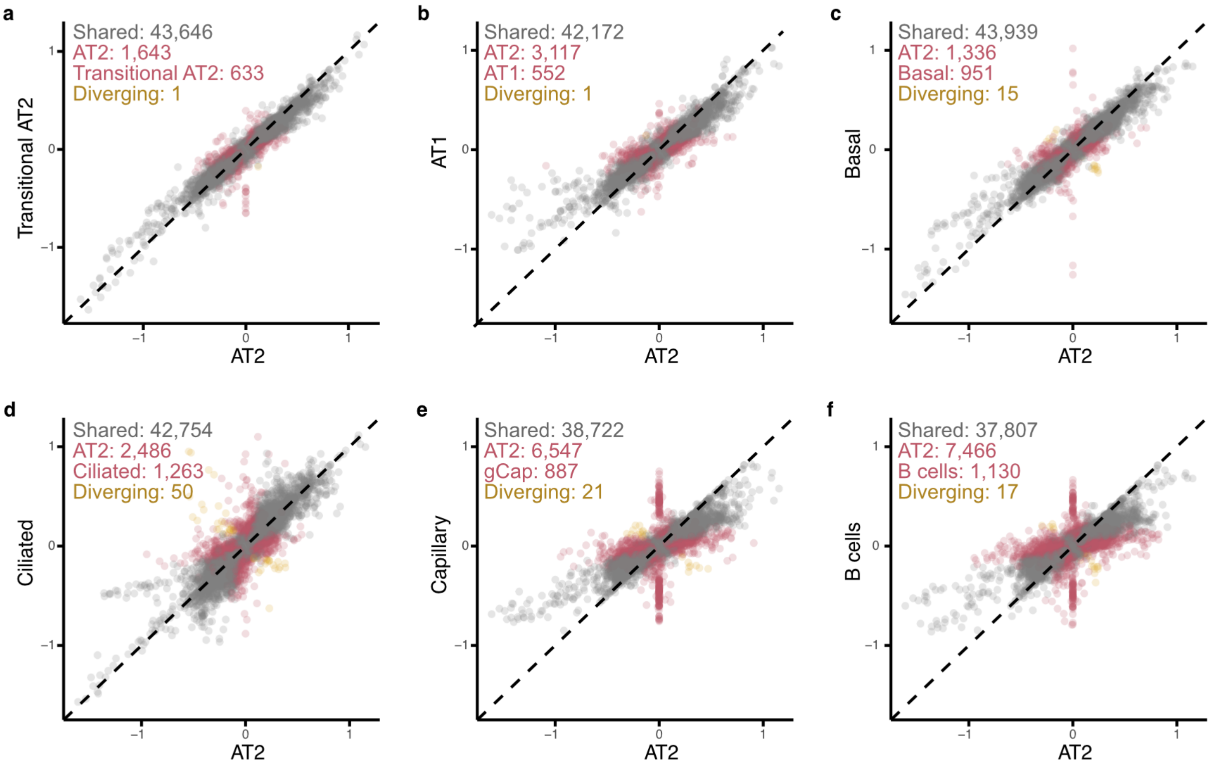


**Fig. S4:** Comparison of mashr estimated effect sizes for top eQTL between AT2 and other cell types. The number of top eQTL that are significant and in the same direction (shared), significant only in AT2, significant only in the comparison cell type, or significant in both but with an opposite direction of effect (diverging) are reported.

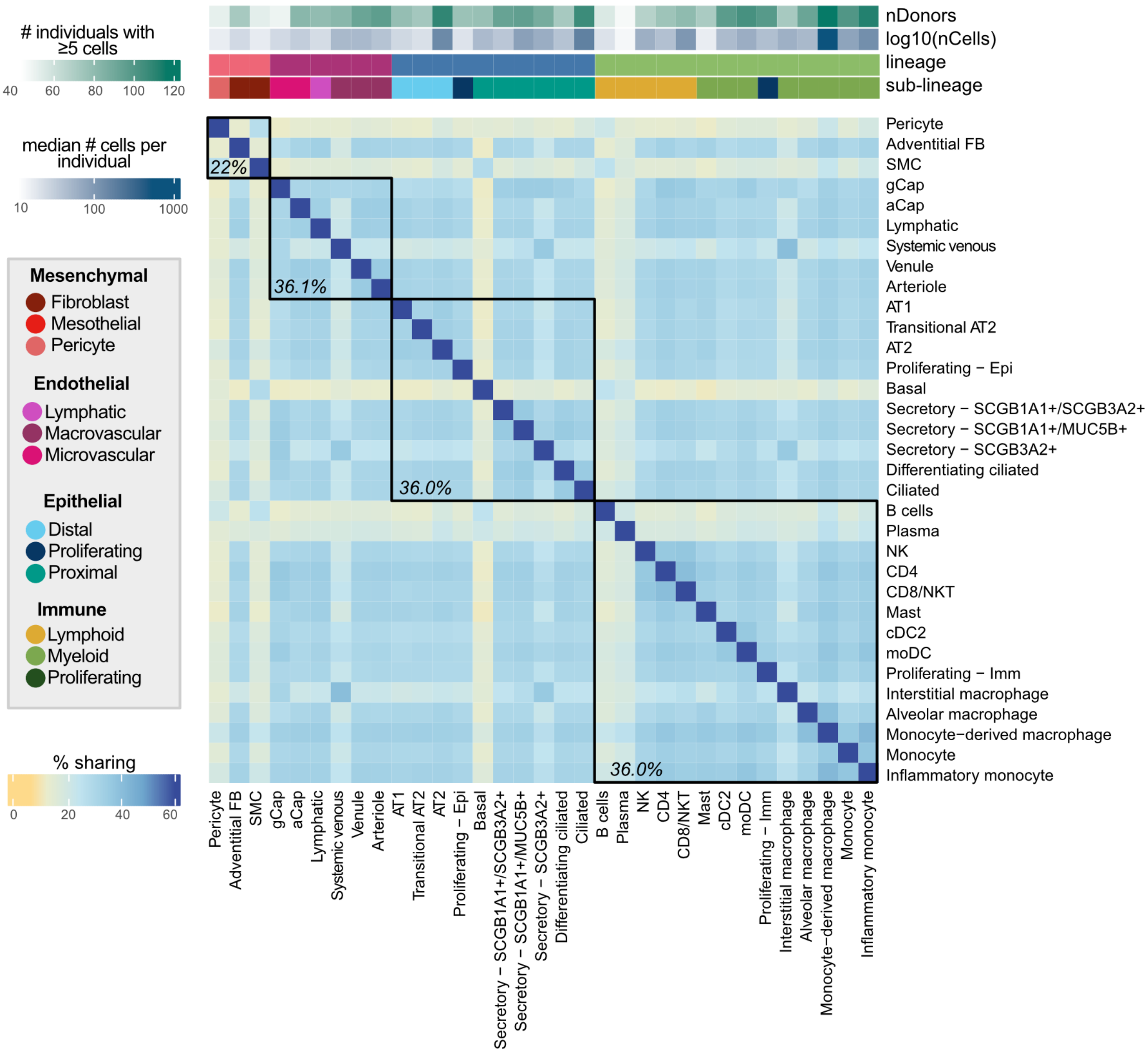


**Fig. S5:** Percent of top int-eQTL that are shared between two cell types. Top eQTL are considered shared if they are significant in both cell types (local false sign rate ≤ 0.1) and the mashr estimated effect size is within a factor of 0.5. Cell types are annotated above by lineage, sublineage, the number of individuals with ≥5 cells, and the median number of cells per individual for that cell type. Median pairwise percent sharing per lineage is shown in black.

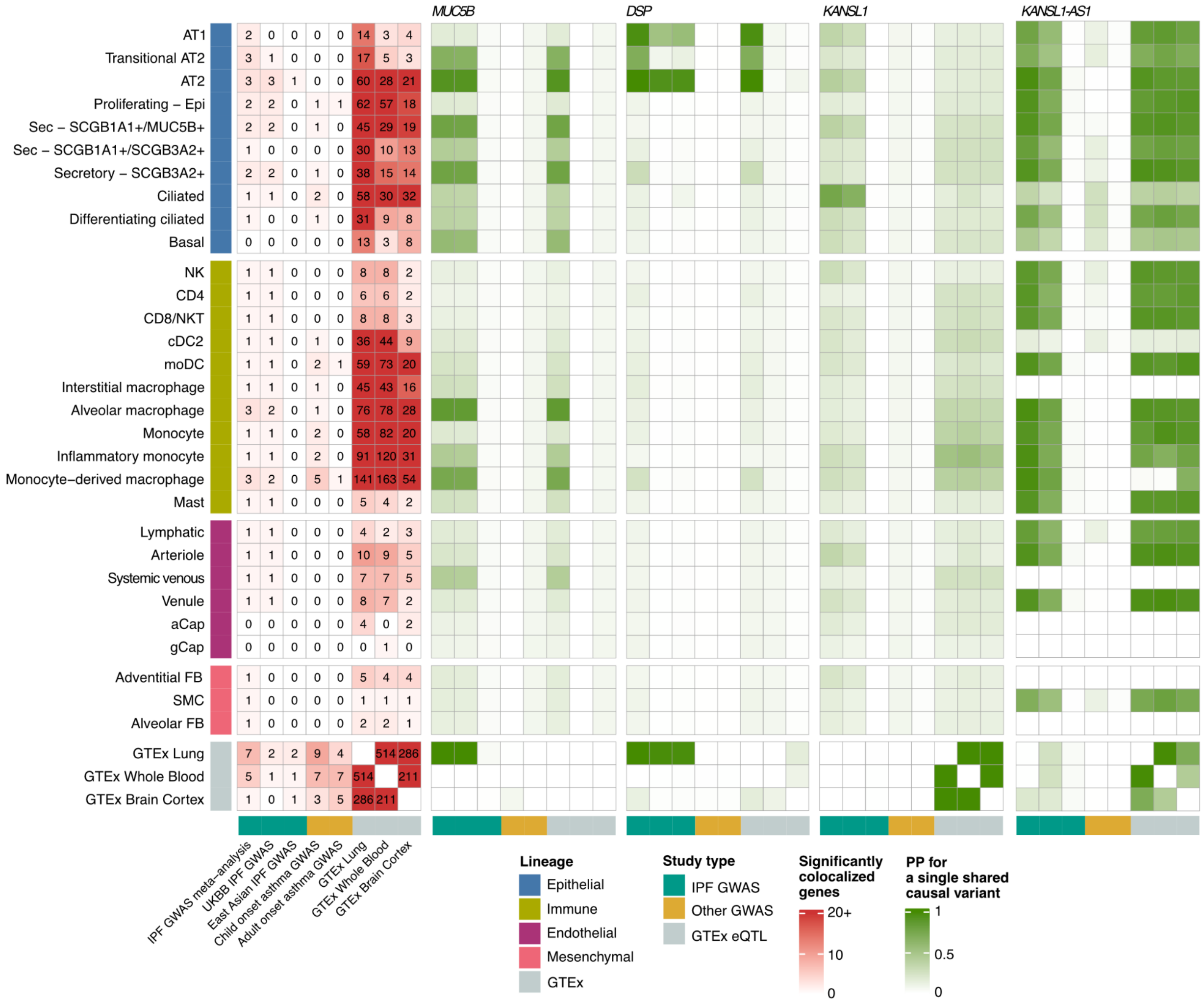


**Fig. S6:** Patterns of colocalization between cell type-eQTL, bulk-eQTL, and lung trait GWAS. The left-most heatmap presents the numbers of significantly colocalized genes between the 38 cell types and three GTEx tissues, three IPF GWASs, child and adult-onset asthma GWAS. The green-shaded heatmaps present posterior probabilities for a single shared causal variant between the tested cell types and GWAS for top IPF GWAS implicated genes (*MUC5B*, *DSP*, *KANSL1*, *KANSL1-AS1*).

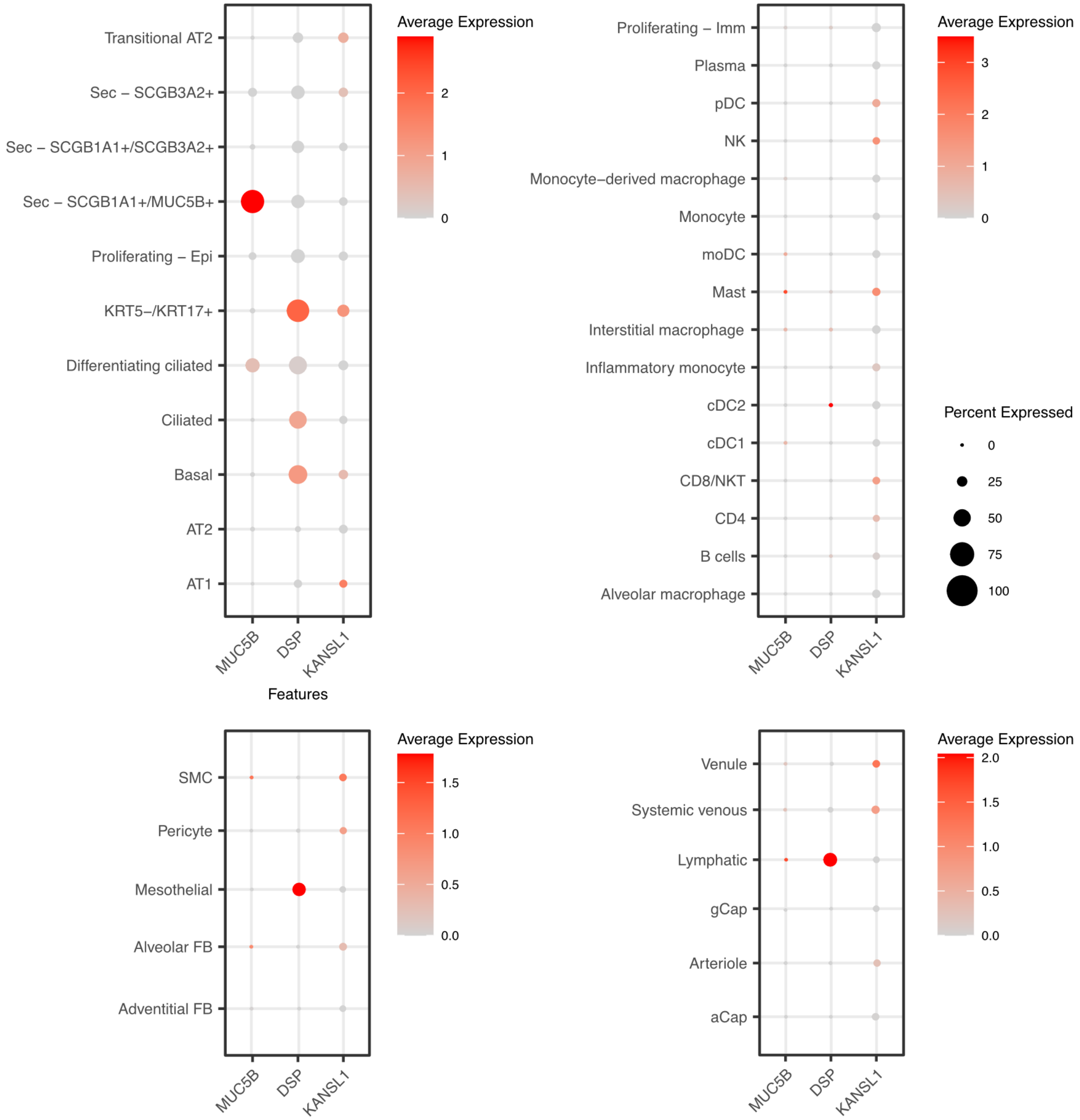


**Fig. S7:** Expression of *MUC5B* across tested cell types.

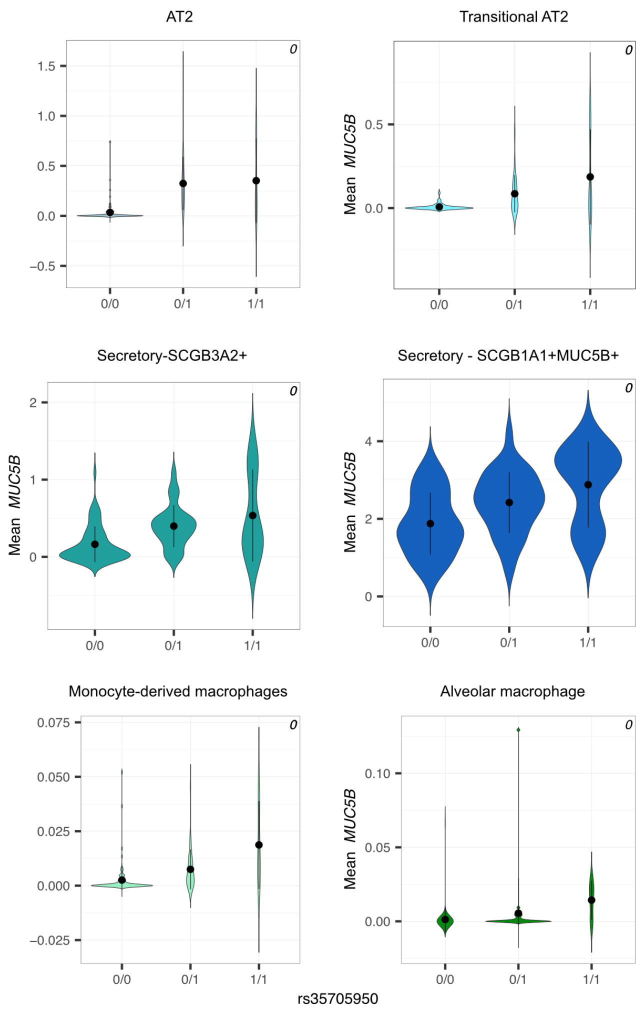


**Fig. S8:** eQTL violinplots of the top GWAS variant for *MUC5B* in cell types in which significant colocalization with GWAS was detected.

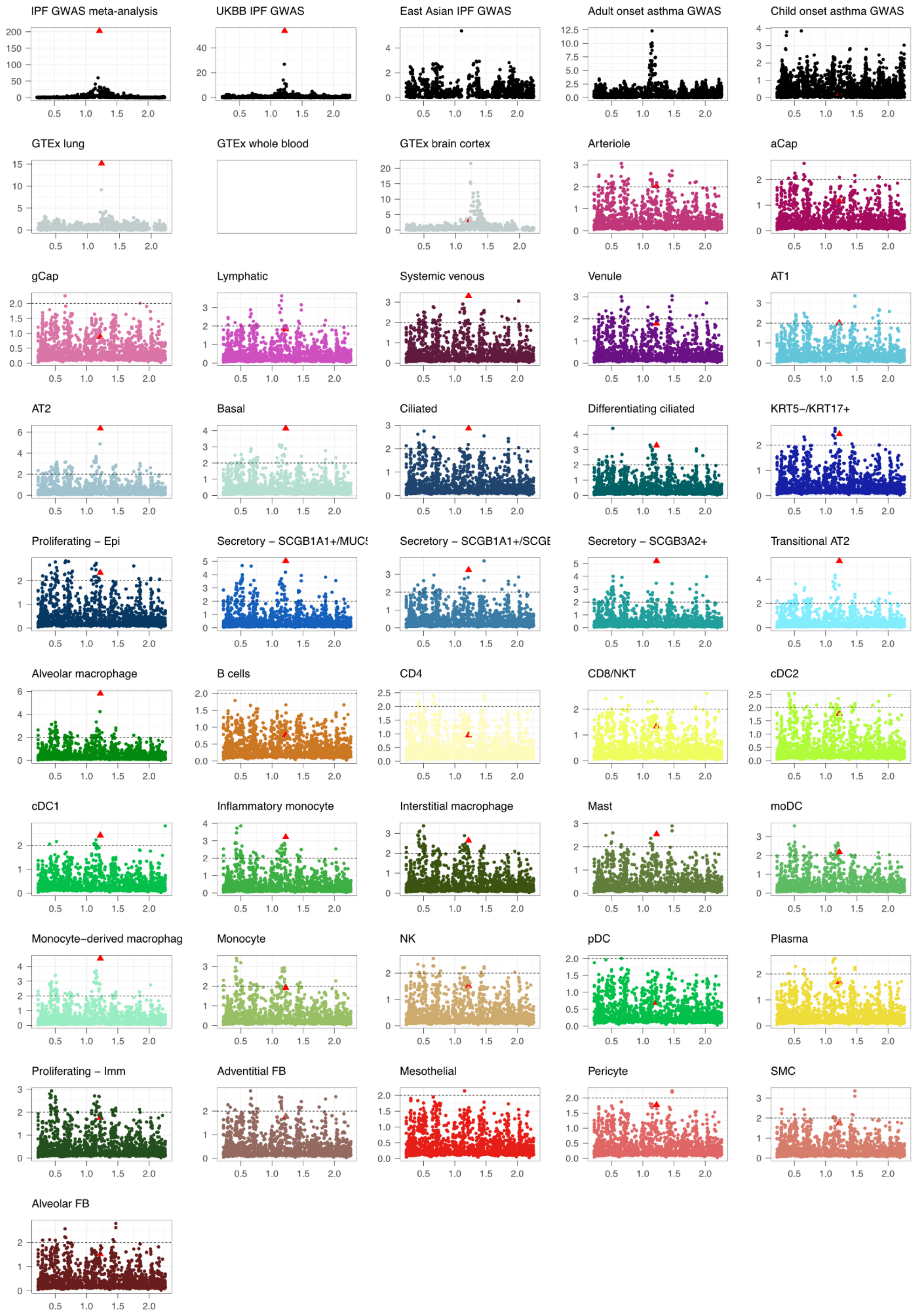


**Fig. S9:** Manhattan plots of -$\log_{10}$ *p*-values or mashR lfsr-values (y-axis) for for *MUC5B* eQTL and GWAS. Basepare positions are indicated in Mb. IPF GWAS top variant is indicated as a red triangle.

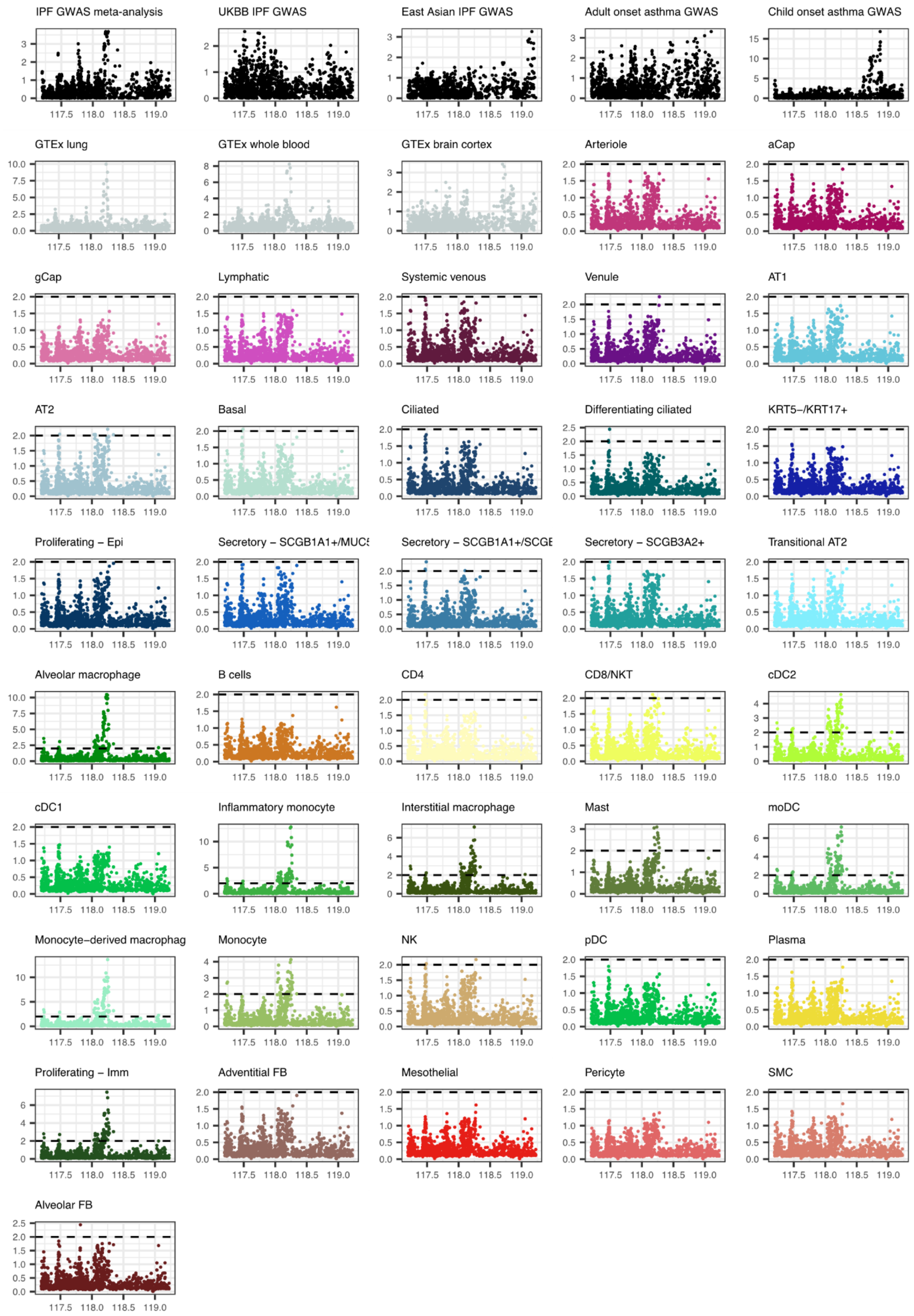


**Fig. S10:** Manhattan plots of -$\log_{10}$ *p*-values or mashR lfsr-values (y-axis) for for *JAML* eQTL and GWAS. Basepare positions are indicated in Mb.

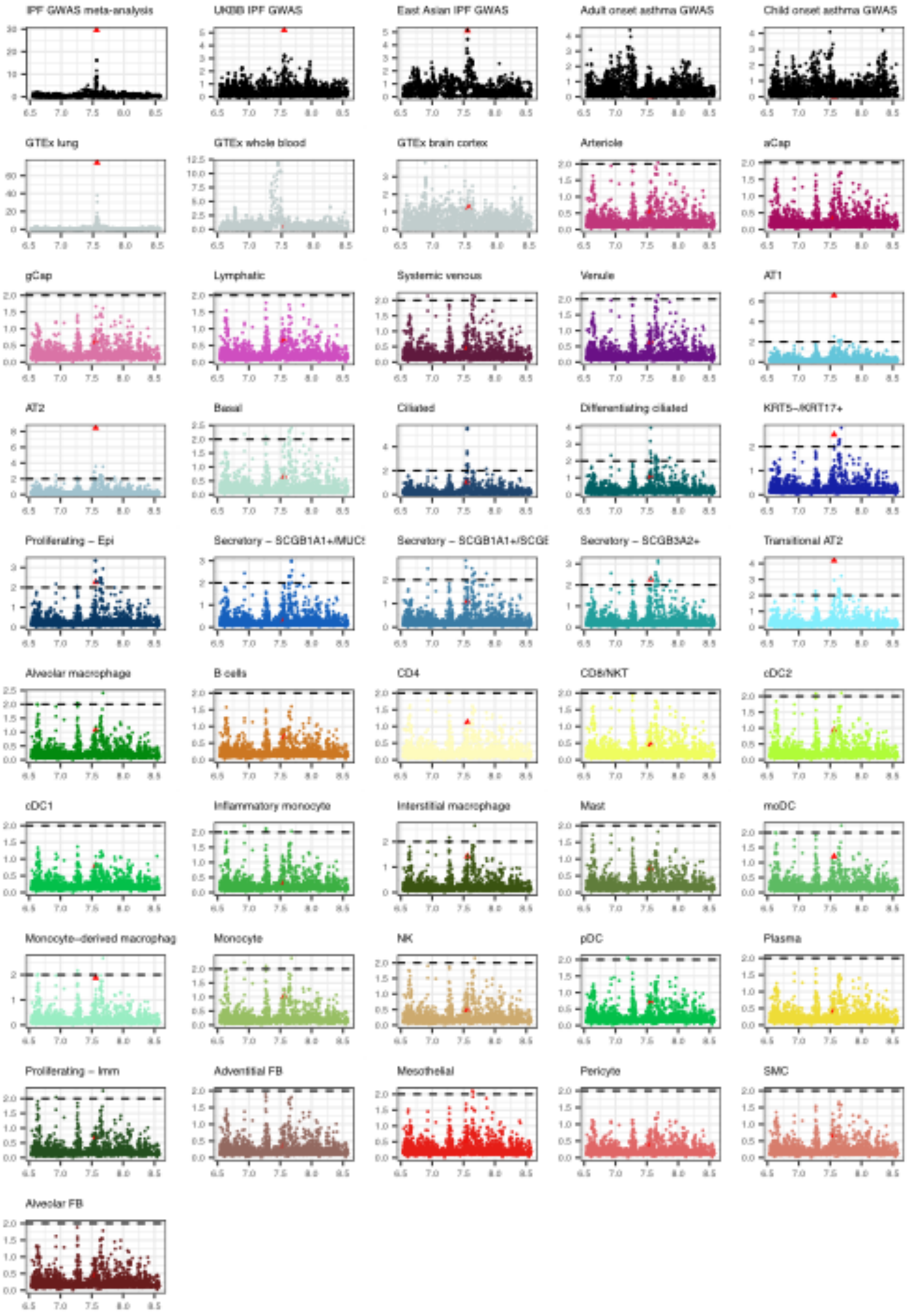


**Fig. S11:** Manhattan plots of -log$_{10}$ *p*-values or mashR lfsr-values (y-axis) for for *DSP* eQTL and GWAS. Basepare positions are indicated in Mb. IPF GWAS top variant is indicated as a red triangle.

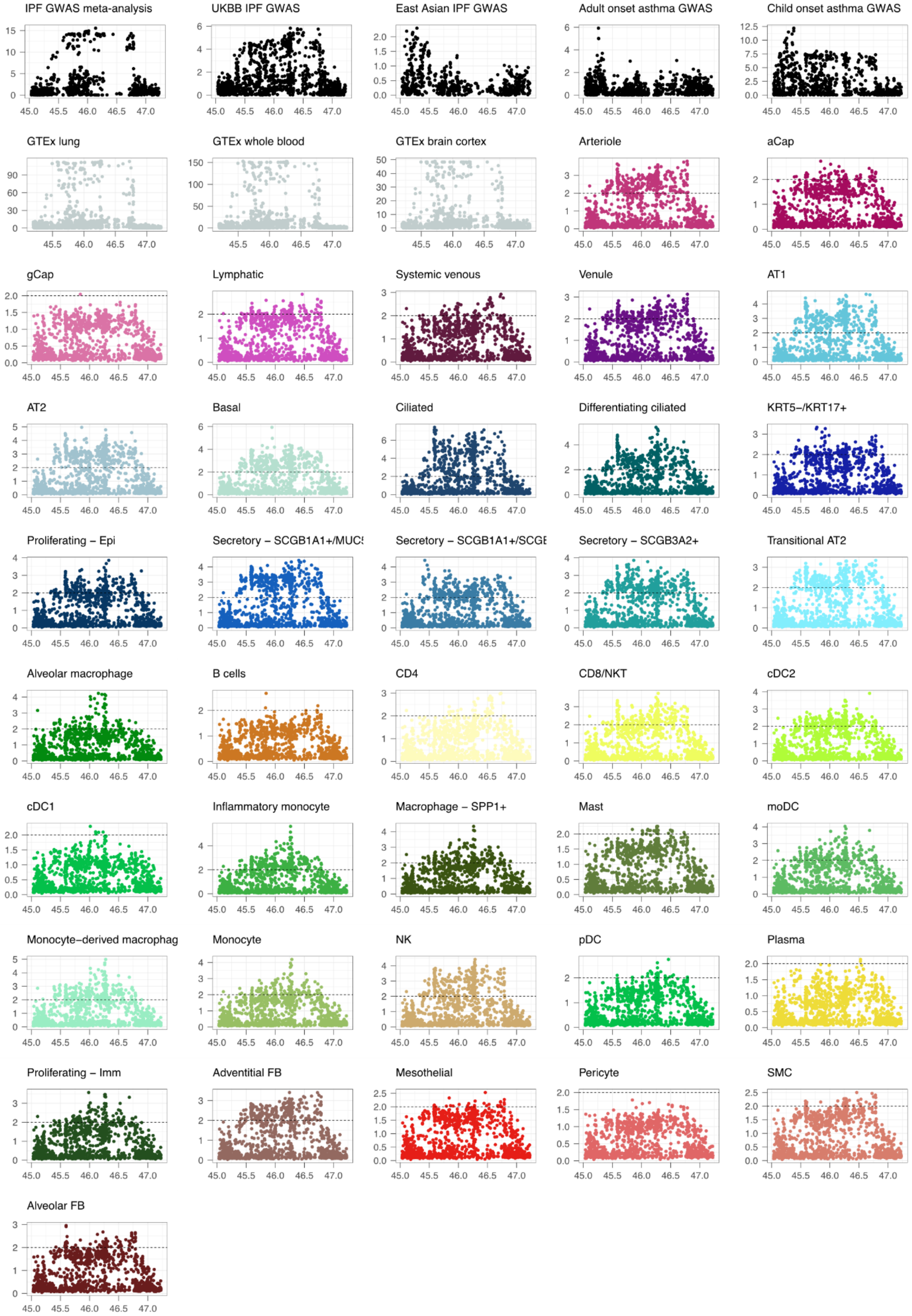


**Fig. S12:** Manhattan plots of -$\log_{10}$ *p*-values or mashR lfsr-values (y-axis) for for *KANSL1* eQTL and GWAS. Basepare positions are indicated in Mb.

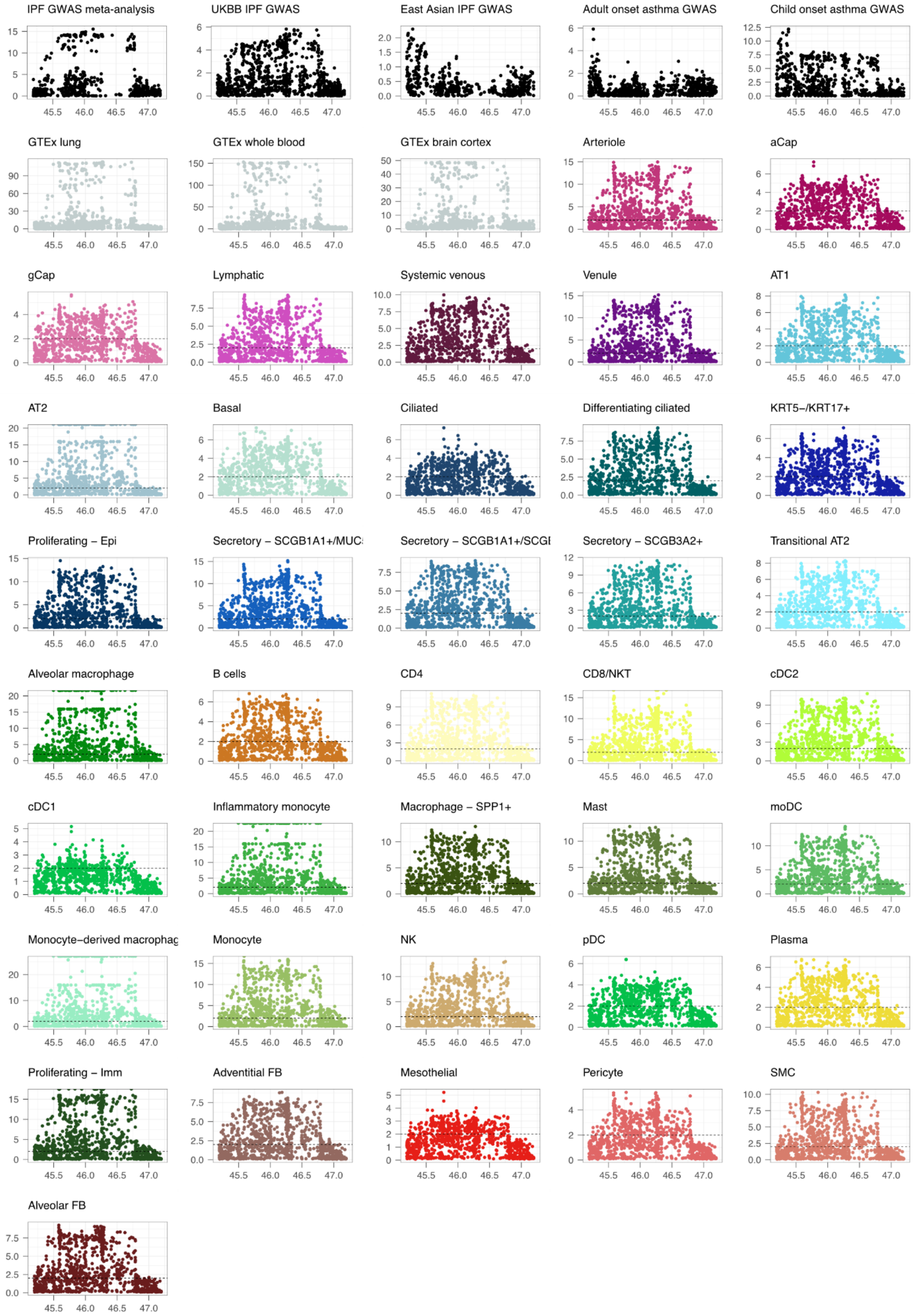


**Fig. S13:** Manhattan plots of -$\log_{10}$ *p*-values or mashR lfsr-values (y-axis) for for *KANSL1-AS1* eQTL and GWAS. Basepare positions are indicated in Mb.

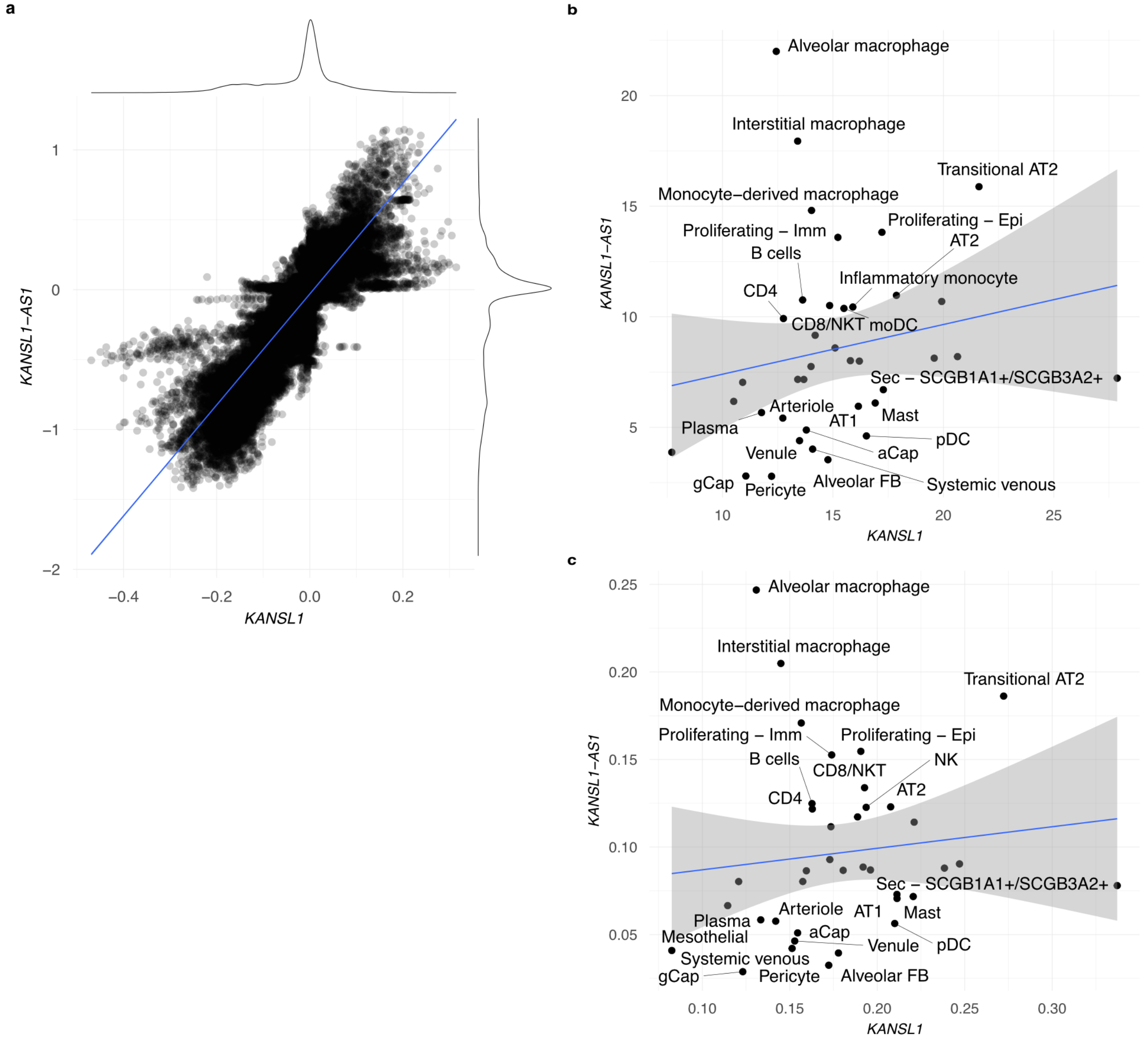


**Fig. S14: a,** *KANSL1* and *KANSL1-AS1* eQTL mashR posteriors. **b,** Proportion of cells expressing and **c,** average expression of *KANSL1* and *KANSL1-AS1* in tested cell types. Outliers are labeled.
